# A Functional Metabolomics Framework to Track Microbiome Drug Metabolism

**DOI:** 10.64898/2026.01.30.702925

**Authors:** Abzer K. Pakkir Shah, Anne Griesshammer, Paolo Stincone, Jarmo-Charles Kalinski, Axel Walter, Mingxun Wang, Lisa Maier, Daniel Petras

## Abstract

Understanding how gut microbes transform drugs, and how this influences microbiome composition and function, remains a key question to better understand the efficacy and side effects of pharmaceuticals. To accelerate the discovery of microbiome-derived drug metabolites, we developed a functional metabolomics workflow that combines the use of synthetic microbial communities (SynComs) with a time-series resolved molecular networking approach and advanced computational metabolite annotation. We demonstrate how this framework can be used to illuminate chemical transformation dynamics in a gut SynCom (Com20) with 50 clinical drugs. Our results highlight a multitude of drug metabolites, including multi-step metabolic cascades, some of which correlated to shifts in microbial taxa, suggesting functional links between microbiome composition and biochemical transformations. Our computational data analysis workflow is publicly available through the GNPS2 ecosystem at chemprop.gnps2.org, which can be used to prioritize biotransformations and other (bio)chemical reactions in various biological and abiotic systems.

## Introduction

The human gastrointestinal (GI) tract hosts an estimated 10^13^ microbes that perform essential functions, including immune modulation, defense against pathogens, and nutrient processing^1–4^. Among these roles, the ability of gut microbes to chemically modify xenobiotics (e.g. pharmaceuticals, food additive, and environmental pollutants) is gaining increasing recognition^3^. These microbial transformations, observed in both humans and animal models, can substantially alter the bioactivity, bioavailability, and toxicity of xenobiotics^5,6^. Unlike the host, which primarily relies on oxidative and conjugative reactions (e.g., via cytochrome P450s) to metabolize xenobiotics, gut microbes predominantly perform hydrolysis and reduction^6^. These transformations often arise from broad substrate promiscuity rather than evolutionary selection. However, identifying the microbial contributors to these transformations remains challenging due to horizontal gene transfer and strain-level variation, and the limited predictability of function from taxonomy alone^3^. Beyond these canonical microbial reactions^7^, recent studies have shown oxidation, demethylation, and conjugative transformations, including amino acid and peptide conjugations, distinct from hepatic Phase II metabolism^8–11^. Importantly, the microbiome function can be highly personalized, shaped by environmental factors, including diet and medication, and can influence how drugs are processed or tolerated^4,12,13^.

In parallel, there is increasing awareness of how drugs can directly affect the microbiome itself. Antibiotics, while essential for fighting infections, are known to disrupt commensal bacterial populations, leading to dysbiosis and associated health consequences such as *Clostridium difficile* infections, metabolic dysfunction, allergic, and inflammatory disorders^14–18^. Despite growing recognition of these effects, the activity spectrum of different antibiotic classes on gut bacteria remains poorly characterized, largely due to technical challenges in culturing anaerobic species and limited susceptibility data for many common or clinically relevant gut microbes^17,19^. Beyond antibiotics, recent studies have demonstrated that non-antibiotic drugs, including antipsychotics, proton pump inhibitors (PPIs), and NSAIDs, can also inhibit the growth of gut microbes. Maier et al.^20^ screened over 1,000 marketed drugs against 40 representative gut strains and found that nearly a quarter (24% of these human-targeted compounds) inhibited at least one strain. Findings from human cohort studies^21–24^ similarly identify several non-antibiotic drugs as risk factors for microbiome disruption and infection similar to antibiotics.

Enzymatic activity from gut microbes can profoundly alter the fate of xenobiotics, enhancing or reducing their activity, generating toxic byproducts, altering stability, affecting absorption, or accelerating elimination^9^. Despite these consequences, the reaction dynamics of such microbiome-mediated chemical transformations are rarely studied. In time-series metabolomics datasets, which can contain thousands of features, it is not only the parent drug (i.e., its precursor ion [M+H]^+^) and its final products that are relevant, but also the transient intermediates formed along the way. Inferring directionality is essential for reconstructing transformation pathways, distinguishing precursors from products, and identifying intermediates that may be bioactive or toxic^9^.

To address the complexity of microbiome-drug interactions, recent efforts have leveraged multi-omics approaches^24–35^, including non-targeted LC-MS and LC-MS/MS to systematically profile the depletion of drugs by gut bacterial strains. However, most methods are tailored for known metabolic reactions and targeted enzyme assays and are not readily applicable for data-driven discovery of microbiome-associated transformations in complex metabolomics datasets.

Molecular networking offers a powerful data analysis strategy to organize and annotate non-targeted MS/MS data, which allows for the rapid prioritization of putative analogs and transformation products^36,37^. While several software tools have been developed to annotate chemical transformations in metabolomics data, most focus on structure prediction rather than transformation dynamics. For example, BioTransformer uses curated reaction rules and machine learning to predict phase I and II metabolism products^38^, and MetWork integrates predicted enzymatic reactions with spectral simulation to suggest candidate structures within molecular networks^39^. While non-targeted metabolomics offer a powerful tool to capture microbial chemical activity, most workflows do not infer transformation directionality or abundance changes over full time series.

Emerging functional metabolomics tools expand the comprehensive measurement of metabolites in a biological system by linking them to specific biological phenotypes or molecular interactions^40^. Such strategies typically integrate orthogonal approaches such as perturbation, including gene knock-outs or knock-downs, protein binding, and bioactivity screens^41–43^. To address the challenge of prioritizing microbiome derived drug metabolism, we developed a functional metabolomics framework that integrates the use of synthetic microbial communities (SynComs), 16S amplicon sequencing, non-targeted metabolomics, and a correlation-based molecular networking workflow (ChemProp2), as well as repository-scale metabolite contextualization. By resolving transformation directionality and mapping multi-step cascades, ChemProp2 enables the identification of microbial drug metabolism and the linking of specific transformations to shifts in microbial community composition.

Here, we highlight the application of our workflow to investigate microbial metabolism of 50 clinically relevant drugs by a gut SynCom (Com20), uncovering drug-specific transformation patterns associated with microbial shifts. To further contextualize unannotated xenobiotic metabolites, we incorporated repository-scale searches with the FASST software tool^44^, enabling a broader interpretation of their biological occurrence and relevance. This integrated approach supports the large-scale matching and contextualization of microbiome-mediated chemical transformations to publicly available non-targeted metabolomics studies. Together these tools provide a framework to better understand microbiome drug metabolisms, which could be leveraged in future drug discovery and personalized medicine approaches.

## Results

### ChemProp Infers Direction of Microbiome-Driven Drug Metabolism

To evaluate drug biotransformation within Com20, we incubated the SynCom in parallel with 50 drugs, followed by non-targeted LC-MS/MS based metabolomics (**Figure 1**). We then applied the ChemProp computational pipeline which computes ratio-based scores between feature pairs in molecular networks to infer putative reaction directionality and is particularly suited for endpoint designs^45^ (**Figure 1**). To address dynamic changes across multiple timepoints, we developed ChemProp2, which integrates full time-series information using correlation-based scoring (**Figure 4A**) and provides an empirical false discovery rate (FDR) procedure via a decoy-based strategy^46,47^. These scores range from -1 to +1, where the magnitude reflects strength and the sign indicates the inferred direction of potential transformation. By comparing scores derived from randomized (decoy) feature tables to real networks, users can apply thresholds (e.g., 1%, 5%, 10% FDR) to prioritize high-confidence transformations. Importantly, ChemProp2 allows for analysis of primary edges (i.e., direct connections to the drug’s precursor ion node) and distal cascade-level relationships, capturing subtle or multi-step transformation events.

**Figure 1:**
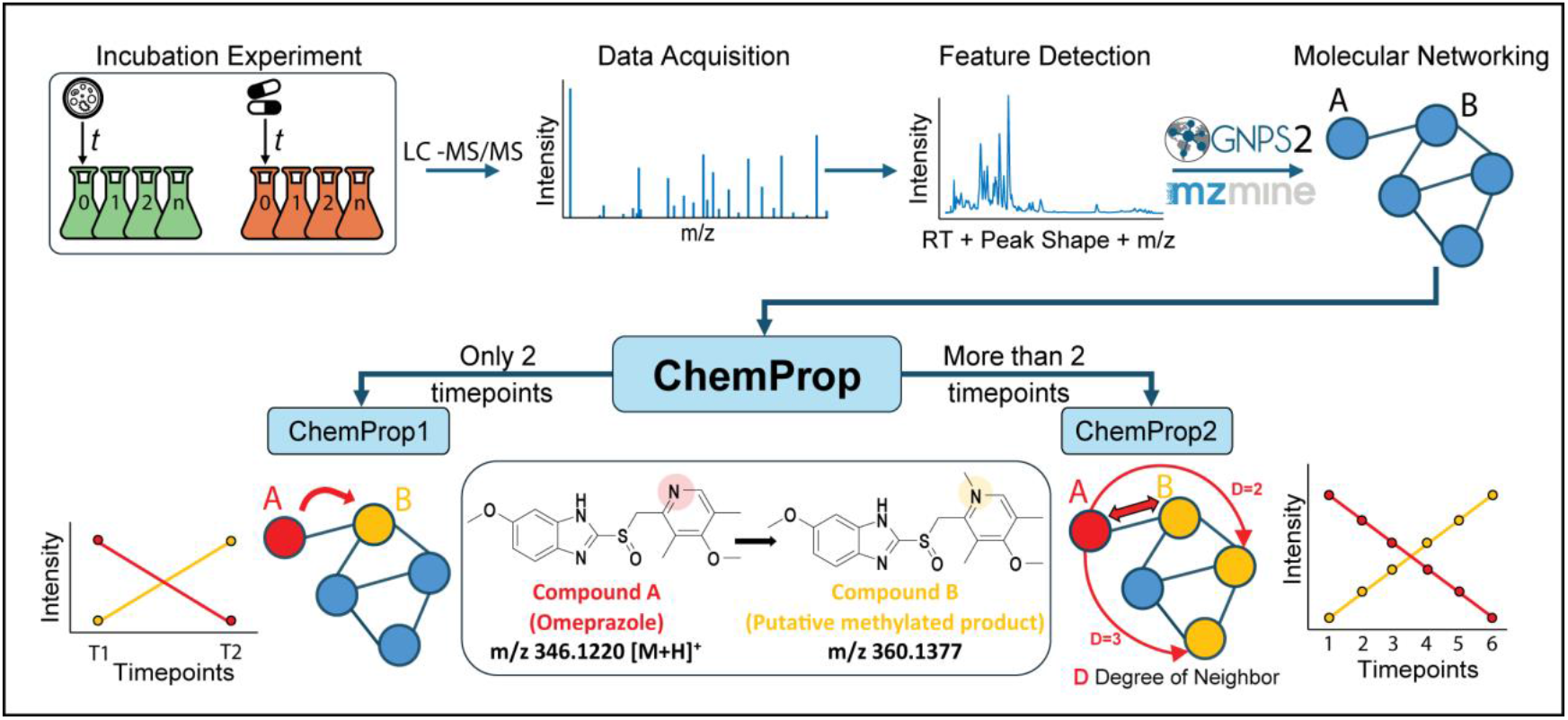
Schematic overview of the ChemProp workflow for detecting directional biotransformations. Starting from a microbial incubation experiment, non-targeted LC-MS/MS data are collected and processed using MZmine for feature detection. Feature-based molecular networking (FBMN) is then performed via GNPS2 to organize structurally related features. ChemProp scoring is applied to the resulting network: ChemProp1 supports two-timepoint designs, while ChemProp2 analyzes multi-timepoint data using correlation-based scoring. Both modules prioritize network edges based on anti-correlated intensity patterns to infer directional transformations. The middle panel shows the molecular structures of Omeprazole and a putative methylated metabolite (*m/z* 360.1377), represented here as an example from our dataset. (The ChemProp web application is available at https://chemprop.gnps2.org).

To make ChemProp broadly accessible, we developed a web application (https://chemprop.gnps2.org/) as part of the metabo-apps^48^ in the GNPS2 ecosystem. The app allows users to upload feature quantification table, metadata, and molecular network edge files. It returns scored edge table with directional information as CSV file, as well as GraphML files for network visualization. Users can interactively filter by score range, focus on specific subnetworks, and inspect corresponding intensity trends for selected feature pairs (see Supplementary Methods). A summary of the overall approach and decision points for using ChemProp1 versus ChemProp2 is illustrated in **Figure 1**. By integrating ChemProp2 with Feature-Based Molecular Networking (FBMN), non-targeted metabolomics data can be coupled with time-resolved scoring to suggest potential biological directionality (e.g., drug to product) within otherwise undirected molecular networks.

### Initial Screening and Drug Prioritization

We employed Com20, a previously established synthetic gut bacterial community comprising 20 commensal bacterial strains spanning six phyla, 11 families, and 17 genera, collectively encoding around 61 % of the metabolic pathways found in a healthy human gut microbiome^24^. The community showed stable and reproducible growth in gut-mimetic mGAM medium under anaerobic conditions^24,49^. To investigate microbiome-mediated biotransformation of small molecules, Com20 was incubated with 50 clinically relevant drugs under anaerobic conditions. Samples were collected at T = 0 h (immediately after drug exposure) and T = 2 h (after incubation) and analyzed using non-targeted LC-MS/MS (**Figure 2A-B**). This two-timepoint setup, referred to as the ChemProp1 model, served as a preliminary experiment to identify drugs exhibiting measurable transformation patterns (See example in **Figure 2C**).

**Figure 2:**
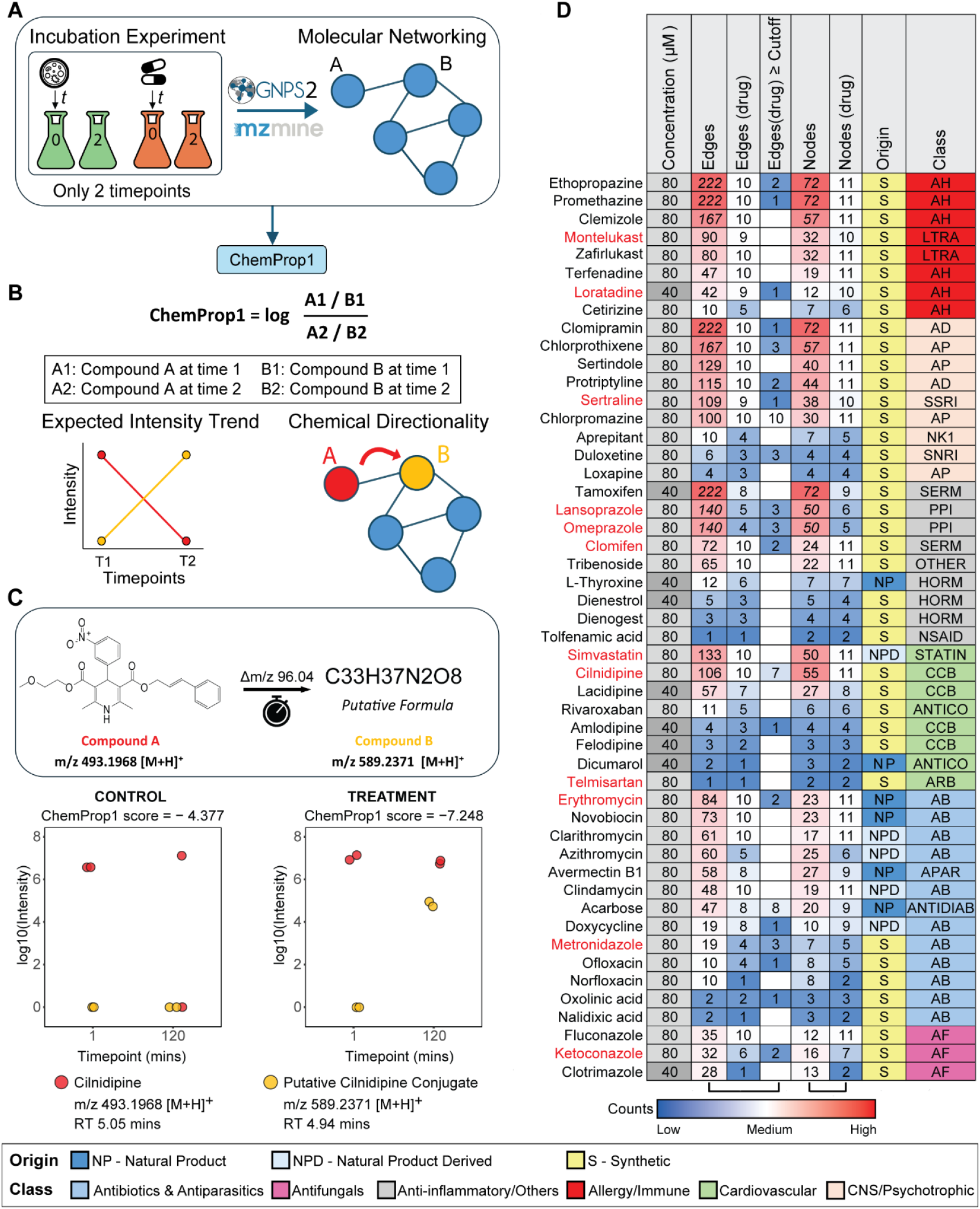
Initial Screening of 50 Drugs Using ChemProp1. (A) Experimental overview: 50 clinical drugs were incubated with the Com20 synthetic gut community under anaerobic conditions and sampled at two timepoints (T0, T2; 2 h apart). Extracts were analyzed by untargeted LC-MS/MS, processed through FBMN, and scored using ChemProp1. (B) ChemProp1 scoring formula based on log-ratio intensity changes between timepoints to infer precursor-product directionality. (C) Example transformation: Cilnidipine (*m/z* 493.1968) to a putative downstream product (*m/z* 589.2371; *Δm/z* = 96.0403). Scatter plots show log-intensity trends in Com20 vs abiotic control. (D) Heatmap summary of ChemProp1 results across all 50 drugs. For each drug, we report total nodes and edges in the main [M+H]^+^ cluster, first-degree connections, and edges above the ChemProp1 threshold (score ≥ 1). Color scale represents edge counts (red = high, white = mid, blue = low/none). Columns additionally indicate drug origin (NP, NP-derived, synthetic) and drug class. Full class abbreviations are listed in Supplementary Table 1. Drugs marked in red were selected for ChemProp2. Italicized values (e.g., Omeprazole/Lansoprazole: 140 edges) denote that the drugs belonged to the same cluster.

All 50 drugs had confidently detected precursor [M+H]^+^ ions and surrounding network connections in FBMN. However, the number of direct edges connected to each precursor ion did not necessarily correlate with transformation likelihood as captured by the ChemProp1 scores. **Figure 2D** summarizes the ChemProp1 screen, showing the number of network connections per drug and how many suggested potential biotransformation for each drug. From this, we selected 12 drugs for time-resolved ChemProp2 analysis. Nine drugs (Cilnidipine, Clomifen, Erythromycin, Ketoconazole, Lansoprazole, Loratadine, Metronidazole, Omeprazole, and Sertraline) showed at least a two-fold change between 0 h and 2 h, suggesting potential microbial conversion and making them strong candidates for follow-up. Among these, Erythromycin and Metronidazole are antibiotics, while the remaining compounds represent non-antibiotic drugs spanning cardiovascular, hormonal, antifungal, gastrointestinal, allergy, and CNS indications. To evaluate ChemProp2’s sensitivity in low-signal or borderline cases, we included three non-antibiotic drugs, Montelukast, Simvastatin, and Telmisartan, that did not pass the ChemProp1 cutoff but retained molecular network connectivity. Together, the selected compounds span six major therapeutic categories represented in **Figure 2D**.

### Time-Resolved Analysis of Drug Biotransformations

To investigate microbiome-mediated transformations in greater detail, we decided to use the 12 prioritized drugs and incubated them with Com20 across a 0-8 h time course, sampled at nine timepoints (T0-T8, hourly) **(Figure 3A)**. After LC-MS/MS analysis, the resulting networking and ChemProp2 results showed temporal correlations between connected features, enabling directional inference of precursor-product relationships.

**Figure 3:**
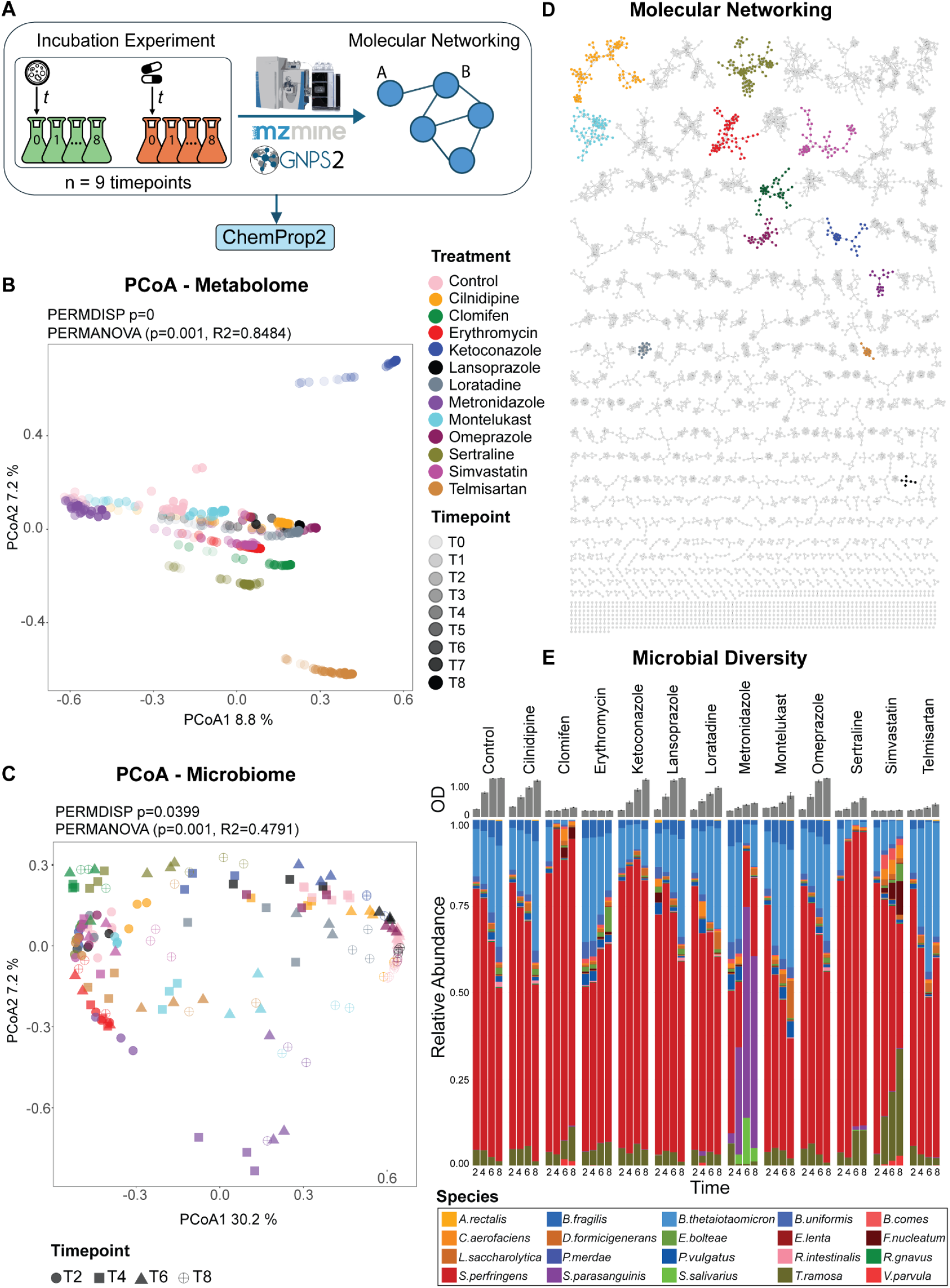
Time-Resolved Multi-Omics Profiling of Microbiome-Drug Interactions. (A) Experimental design: 12 drugs were incubated with the Com20 microbial community and sampled at nine timepoints (T0-T8) in triplicates. Samples were analyzed by non-targeted LC-MS/MS and 16S rRNA sequencing. Metabolomics data were processed via FBMN and subjected to ChemProp2 scoring to identify directional biotransformations. (B) Principal Coordinates Analysis (PCoA) of metabolomics profiles based on Bray-Curtis dissimilarity across all timepoints (T0-T8), showing distinct trajectories for each drug treatment. (C) PCoA of OD-corrected microbial community composition based on 16S rRNA sequencing at four timepoints (T2, T4, T6, T8) using Bray-Curtis dissimilarity, highlighting treatment-specific differences. (D) FBMN molecular network of all detected features, with the [M+H]^+^ ion cluster of each of the 12 drugs highlighted in distinct colors. (E) Stacked bar plots of microbial species-level composition at the four timepoints under drug treatment versus control conditions, illustrating dynamic shifts in community structure over time. For each timepoint, species-level relative abundances from the biological replicates were combined and normalized such that each bar sums to one, illustrating temporal shifts in community structure.

Principal Coordinates Analysis (PCoA) using Bray-Curtis dissimilarity (**Figure 3B**) was performed on the metabolomics dataset (8,055 features, reduced to 5,321 after background removal, imputation and TIC normalization). PCoA showed clear separation of metabolic profiles across treatments (PERMANOVA p = 0.001, R^2^ = 0.85; PERMDISP p = 0). Drug-specific clustering in PCoA was most pronounced for Ketoconazole and Telmisartan, suggesting divergent metabolic profiles over the 0-8 h time course. For Telmisartan, this divergence became apparent in the extended multi-timepoint dataset, whereas ChemProp1 captured no transformation signal at the 0-2 h range. Notably, this separation reflects shifts in global metabolite profiles captured by Bray-Curtis PCoA and is independent of cascade depth or network size, as shown by the varying FBMN cluster sizes in **Figure 3D**. While angiotensin receptor blockers have been reported to undergo microbiome-associated transformation^50^ and alter microbial composition in vivo^51,52^, the specific features driving Telmisartan’s separation here remain unclear and warrant further investigation. The result highlights that drugs with low initial transformation signals can nonetheless produce distinct metabolic trajectories when monitored over time.

At the community level, PCoA on OD-corrected 16S rRNA data (**Figure 3C**) across timepoints T2-T8 revealed modest treatment-based separation (PERMANOVA p = 0.001, R^2^ = 0.48). Variation along PCo1 reflected temporal progression, with T2 samples at one end (**SI Figure 2**), while PCo2 showed a weaker drug-associated signal, most notably for Metronidazole. Superimposed on this temporal gradient, several treatments (e.g., Erythromycin, Clomifen, Sertraline, Simvastatin, Telmisartan) clustered in the upper-left region of PCo1 and Pco2, whereas DMSO control and DMSO-like profiles (Omeprazole, Lansoprazole, Cilnidipine) were positioned toward the upper-right region of PCo1 and Pco2. Individual PCoA plots for each drug treatment and the DMSO control (**SI Figures 3-4**) further supported these trends: most treatments displayed clear temporal separation, with earlier timepoints (T2-T4) consistently diverging from later samples. In contrast, metabolomic PCoA trajectories (**Figure 3B**) showed more complex temporal patterns, while overall progression was evident, a few samples (typically at T0 or T4-T5) strongly influenced variation along PCo1.

Metronidazole, an antibiotic widely used against *Clostridioides difficile*, produced the most distinct taxonomic profile, consistent with its antimicrobial activity^17,53^. In Com20, *S. perfringens* is a key member for community stability^24^; its reduction **(Figure 3E)** therefore led to pronounced community restructuring^54^. As *S. perfringens* declined, lower-abundance commensals (*S. parasanguinis, S. salivarius*) increased, aligning with reports of Metronidazole tolerance or resistance^55–57^. The strong response observed here is thus expected when a keystone taxon is targeted. In addition, *E. coli* and *Streptococcus* overgrowth following Metronidazole exposure has been reported previously^58,59^, supporting the patterns observed in our system.

Similar effects were observed for Simvastatin, a statin, with increases in *T. ramosa* and *F. nucleatum* (**Figure 3E**). *T. ramosa* responded to several drugs but increased most consistently under Simvastatin treatment, whereas *F. nucleatum* enrichment appeared unique to this condition (**SI Figure 5**). Previous work reported *E. lenta* and *B. thetaiotaomicron* as Simvastatin-sensitive species showing growth inhibition and transcriptomic signatures of membrane remodeling or drug efflux^60^. Consistently, we observed a decrease in *B. thetaiotaomicron* abundance over time (**Figure 3E**).

To assess overall taxon involvement across different drug treatments, mean OD-corrected 16S rRNA abundances were visualized as heatmaps for each time point (**SI Figure 5**). Most taxa showed low but detectable abundances across treatments, underscoring the broad participation of the community in drug metabolism. This pattern is consistent with earlier reports that many drugs can be transformed by multiple taxa across phyla^50^. Together, these findings suggest that while metabolic shifts occur rapidly and are highly drug-specific, microbiome composition changes more slowly. Microbial perturbation was evident in both metabolomic and 16S datasets, but the stronger effect size in metabolomics (R^2^ = 0.85 vs. 0.48) underscores its greater sensitivity in detecting early biochemical transformations. These results validate ChemProp2 as a framework for uncovering microbiome-mediated drug metabolism and highlight the importance of integrated omics approaches.

Across all 12 drugs, the summary metrics revealed substantial variation in how treatments affected metabolome and microbiome dynamics over time (**SI Figure 6**). Metabolome and microbiome trajectory lengths were positively associated (Spearman r = 0.73). Several drugs, including Lansoprazole, Montelukast, Cilnidipine, Omeprazole, and the DMSO control, showed relatively large temporal shifts in both metabolomic and microbiome PCo1 trajectories, whereas Erythromycin and Clomifen exhibited minimal movement in either layer. Metronidazole displayed a distinct pattern: it induced noticeable microbiome restructuring but only minor metabolomic change. (**SI Fig. 6A**). No clear relationship was observed between the number of ChemProp2-predicted transformations and the magnitude of metabolome PCo1 change across treatments **(SI Fig. 6B)**. Biomass trends aligned with the observations in **SI Fig. 6A**, as treatments with larger OD increases generally also showed greater temporal change along PCo1 in both the metabolome (Spearman r = 0.74) and microbiome (Spearman r = 0.97) **(SI Fig. 6C-D)**, suggesting growth effects partly drive these patterns.

Shannon diversity remained stable for most treatments over the 8-hour incubation period (**SI Table 2**), which is expected given the short timeframe, initial dilution, and typical doubling times of gut bacteria. Importantly, PPIs often formed only small or fragmented FBMN clusters. This likely reflects chemical behavior rather than biological absence of transformation, as PPIs such as Omeprazole generate many structurally diverse, low-abundance degradation products with heterogeneous MS/MS fragmentation patterns^61,62^. Together, these results show that drugs can substantially reshape microbial metabolic activity without proportionally altering community structure.

### Cascade Scoring Reveals Multi-Step Biotransformations

An inherent bottleneck of modified-cosine scoring for molecular networking is that it primarily links metabolites that differ by a single structural modification, as the algorithm compares MS/MS similarity while accounting only for one mass shift (given by the precursor mass difference). Consequently, only one-step precursor-product relationships are typically connected. To move beyond these direct edges, we implemented a cascade scoring approach in ChemProp2 that traverses the molecular network to identify multi-step connections from each parent drug node (**Figure 4A-B**). This is particularly useful for multi-step biotransformations, in which the intermediates have short half-lives and are only present in minor amounts. Starting from 115 first neighbor edges (D1), cascade expansion added 549 additional links to the 14 parent drug nodes, yielding a total of 664 drug-associated edges, a ∼5.8-fold increase compared to D1 alone. Many distal features displayed anti-correlated intensity profiles with their parent drugs, suggestive of sequential metabolic turnover. **Figure 4C** summarizes the distribution of ChemProp2 scores by Δ*m/z*, highlighting recurring shifts corresponding to common modifications such as methylation (+14.02 Da) and oxidation (+16.00 Da).

**Figure 4:**
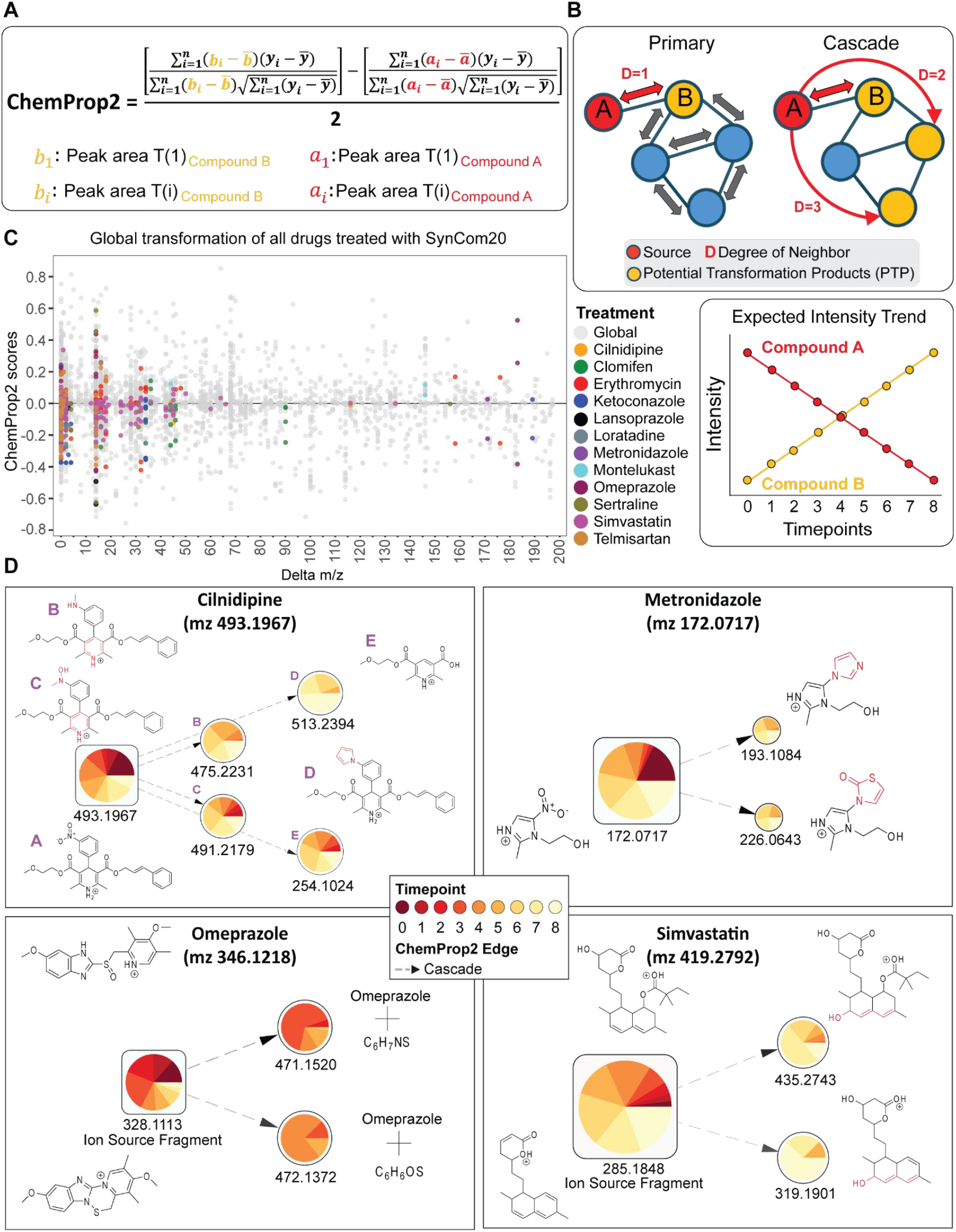
ChemProp2 Scoring Framework and Application to Drug Networks. (A) ChemProp2 scoring formula based on Pearson correlation, the default metric used to quantify directional relationships between feature pairs across multiple timepoints. (B) Conceptual illustration of primary edges (direct drug-feature connections) versus cascade edges (multi-step downstream connections) within a molecular network, and expected intensity trends associated with putative biotransformations are shown. (C) Global ChemProp2 score distribution for all 12 drugs. The x-axis shows observed *m/z* differences between feature pairs, corresponding to potential chemical modifications (e.g., +14 Da for methylation), while the y-axis shows ChemProp2 correlation scores (-1 to +1). Each drug’s primary-edge scores are color-coded, and all other pairwise scores in the network are shown in gray. (D) Molecular subnetworks for four representative drugs (Cilnidipine, Metronidazole, Omeprazole, and Simvastatin), with highlighted putative transformation products (PTPs) and proposed structures.

To investigate the cascade scoring utility, we selected four representative drugs: Cilnidipine, Metronidazole, Omeprazole, and Simvastatin, each exhibited strong treatment-specific ChemProp2 scores not observed in abiotic controls. Although the remaining eight drugs also showed ChemProp2 scores, these were comparable between treatment and abiotic conditions and were therefore excluded. The selected drugs uniquely displayed treatment-specific features, suggesting biologically driven microbial biotransformation.

Cilnidipine (*m/z* 493.1967 [M+H]^+^), a dihydropyridine calcium channel blocker^63,64^, is known to be metabolized hepatically via CYP3A to three major products: dehydrogenated (aromatized) form, a demethylated side-chain product, and a combined dehydrogenation/demethylation metabolite^65^. In our SynCom experiment, 44 treatment-exclusive features were detected, of which a subset with high ChemProp2 scores and interpretable MS/MS spectra are highlighted in **Figure 4D**. MS/MS spectra of the highlighted features were manually interpreted using exact mass and MS/MS fragments (**SI Figure 7**). In addition, pairwise MS/MS spectral comparisons between each highlighted feature and the corresponding parent drug, generated using the Spectrum Resolver in GNPS2^66^, as well as predicted modification sites from ModiFinder^67^, are provided in **SI Figure 8**. The feature at *m/z* 491.2179 was consistent with dehydrogenation of the dihydropyridine ring together with nitroreduction to a hydroxylamine and *N-*methylation, transformations commonly associated with microbial activity^68,69^; *m/z* 475.2231 likely represents an analogous metabolite with full reduction of the hydroxylamine. Additional features at *m/z* 254.1024 and 513.2394 may correspond to products of nitrobenzene cleavage and nitroreduction followed by addition of C_4_H_4_, respectively. These findings suggest that Cilnidipine can undergo both hepatic-like dehydrogenation and additional microbial nitroreductive transformations consistent with previously described gut bacterial activities^7,70^. Clinical pharmacokinetic studies report Cilnidipine peak plasma concentrations at ∼2 h and elimination half-lives of 3-4 h^71^. In our SynCom experiment, however, the major Cilnidipine-derived features appeared later (5-8 h). Several detected products such as ring dehydrogenation resembled known hepatic metabolites^65^, whereas heavier products lacked hepatic analogs and likely represent microbial-specific transformations. Cascade scoring also revealed additional high-scoring mass differences consistent with demethylation, hydroxylation, or conjugation events^3,7,70^. Collectively, these findings suggest that microbial enzymes within Com20 can reproduce hepatic-like dehydrogenation while introducing distinct nitro-group modifications^7,70^.

Metronidazole (*m/z* 172.0717 [M+H]^+^) is a nitroimidazole antibiotic^72^ metabolized hepatically via hydroxylation and conjugation, with 2-hydroxymetronidazole as the predominant product, whereas microbial metabolism is dominated by nitroreduction^73^. In our dataset, cascade scoring revealed two treatment-specific products at later timepoints (T4-T8): *m/z* 193.1084 (+21 Da), consistent with nitro reduction and possible formation of a nitrogen-containing imidazole derivative, and *m/z* 226.0643 (+54 Da), likely representing a conjugated product with a thiazolone-like moiety. Both metabolites showed strong residual precursor intensity and characteristic neutral loss of the ethoxy chain during fragmentation, indicating structural stability for the putative metabolites (**SI Figure 9**). Corresponding MS/MS spectral pair comparisons with the parent drug, along with predicted modification sites from ModiFinder, are provided in **SI Fig. 10A-B**. Their appearance coincided with enrichment of *Sarcina perfringens*, a known nitroreductase producer^54^, underscoring their potential microbial origin. Although known hepatic metabolites such as N-(2-hydroxyethyl)-oxamic acid and acetamide^72,73^ were absent, canonical nitroreduction products and several new metabolites emerged under microbial conditions at later timepoints (5-8 h). The Metronidazole network also highlighted a limitation of relying solely on spectral connectivity, as several features were attributed to neighboring clusters. Among these disconnected nodes, several with larger mass shifts (+94 to +143 Da) exhibited fragmentation patterns consistent with partial ring cleavage and conjugation with amino acid- or peptide-like moieties^3,7,74^. Such transformations, though previously unreported for Metronidazole, are chemically plausible given the widespread occurrence of bacterial nitroreductases and conjugating enzymes in gut microbes^70,75^. Together, these findings extend current models of Metronidazole activation beyond simple nitroreduction and highlight potential downstream fates of reduced intermediates within microbial communities.

Omeprazole-related features formed three distinct clusters: the protonated parent ion (*m/z* 346.1217), a sodium adduct (*m/z* 368.1039), and a dehydrated in-source fragment (*m/z* 328.1113). The latter structure likely arises from water loss involving the sulfoxide oxygen^76,77^. The parent and sodium-adduct clusters included features also present in abiotic controls, suggesting spontaneous or non-microbial processes. By contrast, the in-source-fragment cluster included treatment-enriched features (*m/z* 471.1520 and 472.1372) connected by strong ChemProp2 scores (**Figure 4D**). These exhibited mass shifts of +125.03 Da and +126.02 Da relative to the parent, corresponding to additions of C_6_H_7_NS and C_6_H_6_OS, respectively. Their MS/MS spectra yielded a dominant fragment at *m/z* 328.1113 but lacked additional diagnostic ions, preventing confident structural assignment (**SI Figure 11**). The corresponding MS/MS spectral comparisons and ModiFinder^67^ predictions are provided in **SI Figure 10C-D**. Beyond the main Omeprazole subnetwork, additional annotated features were detected, including Ufiprazole, Esomeprazole magnesium and multiple adducts. Several further treatment- and abiotic-specific features were observed as isolated nodes or small clusters but were not considered further due to limited network connectivity and insufficient spectral support. ChemProp2 revealed treatment-enriched derivatives at *m/z* 471.1520 and *m/z* 472.1372 that likely represent previously uncharacterized conjugates. The corresponding mass shifts suggest addition of heteroatom-containing moieties, possibly through microbial conjugation reactions. Prominent neutral losses of the added atoms indicate that these mass shifts represent distinct substituents, possibly bound by a single bond to the parent structure. Although analogous additions have not been reported for Omeprazole, sulfur- and nitrogen-based microbial conjugations are well documented^7,70^. These observations raise the possibility that anaerobic gut microbes can mediate conjugative transformations beyond those seen in hepatic metabolism. Moreover, as multiple microbial species are capable of transforming Omeprazole^78–81^, parallel cascades may occur simultaneously and complicate the reconstruction of directionality.

Simvastatin (*m/z* 419.2792, [M+H]^+^), yielded low ChemProp1 scores, but ChemProp2 identified ten features unique to the microbial treatment. These included both singletons and connected subnetworks, with some features more abundant than the parent ion. Most appeared within the first 4 h of incubation. For example, the ion at *m/z* 435.2743, likely corresponds to the known metabolite 3′-hydroxysimvastatin^82^ and was clustered with the major in-source fragment (*m/z* 285.1848). Another feature at *m/z* 319.1901 did not match reported derivative and likely represents an in-source fragment of *m/z* 435.2743 (**SI Figure 12**). The corresponding MS/MS spectral comparisons and ModiFinder predictions are provided in **SI Figure 10E-F**. Simvastatin illustrates how ChemProp2 captures microbial transformation products that were missed by the two-timepoint model. Most features in the Simvastatin cluster appeared within the first four hours, outside the 0-2 h window used in ChemProp1, explaining its minimal initial scores. Cascade expansion further increased ChemProp2’s sensitivity, revealing a range of Simvastatin-related products, including the known metabolite 3’-hydroxysimvastatin (*m/z* 435.27) alongside in-source fragments, adducts, and statin analogs. Telmisartan similarly produced weak ChemProp1 scores but yielded detectable ChemProp2 features suggestive of methylation and charge-state shifts.

While several other drugs also showed cascade-level ChemProp2 scores, their associated features were often present in abiotic controls, suggesting non-biological or ambiguous origins. We therefore focused subsequent analyses on the above-mentioned four representative case studies, which best illustrate how ChemProp2 cascade scoring captures both known and candidate microbial-specific transformations in time-resolved metabolomics data. Compared to the earlier two-timepoint ChemProp1 model, ChemProp2 integrates temporal dynamics across multiple timepoints and applies FDR-based correction, resulting in fewer but more robust edges (**Table S3** and **Figure S13**).

### Contextualization of Drug Metabolites against Public Metabolomics Data

ChemProp2 suggested numerous putative drug metabolites across the SynCom time series. Once such candidate metabolites emerge from temporal analysis, a natural question arises: how broadly do these features appear beyond this experimental system, and which ones merit deeper biological investigation? Because direct experimental validation for every feature is not feasible, an intermediate step is needed to assess whether a metabolite is recurrent, condition-specific, or potentially an artifact.

To provide an additional contextual layer, we searched all prioritized MS/MS spectra using the FASST platform^44^, which enables large-scale spectral similarity searches across public databases (more than 2 billion MS/MS spectra, November 2025) . Although public metadata are not uniformly curated, repository matches offer valuable orthogonal evidence, indicating whether a feature is observed across diverse biological or environmental studies, or instead appears to be unique to the microbiome-drug interaction examined here. In total, 13 databases were searched, including curated spectral libraries (e.g., GNPS Library, MassIVE-KB) and repository-scale datasets spanning MassIVE, Metabolomics Workbench, and Pan-Repository collections (full list in SI section *FASST Inquiry*). Across 13 repositories, 1,063 of 1,202 queried features returned at least one match, spanning 1,670 unique MassIVE datasets (including the six datasets generated in this study, for more details: see SI section *FASST Inquiry* (**SI Table 4**). This broad recurrence suggests that several drug-derived features or structural analogs appear across independent human, animal, plant, or environmental studies.

To prioritize putative transformation products (PTPs), we retained cascade nodes that (i) differed from the parent [M+H]^+^ ion by > 0.5 Da, (ii) were observed in external datasets, and (iii) had non-zero ChemProp2 treatment scores (see SI section *Cross-Dataset Distribution of Cascade Nodes*, **SI Table 5**). This filtering resulted in 78 PTPs, summarized in **Table 1** and visualized in a heatmap (**Figure 5**), which shows their distribution across external dataset categories and contrasts ChemProp2 treatment versus control scores. Smaller Δ*m/z* shifts (e.g., −2.02, +14.02, +15.99 Da) corresponded to chemically intuitive modifications such as hydrogen loss, methylation, or oxidation, whereas larger Δ*m/z* values often lacked clear annotation but gained contextual support from repository matches. Representative examples illustrate how public data strengthen interpretation. A −17.97 Da transformation linked to Cilnidipine appeared in one plant-related and three unclassified repository datasets and showed a ChemProp2 treatment score of 0.31, consistent with a plausible biological transformation.

**Figure 5:**
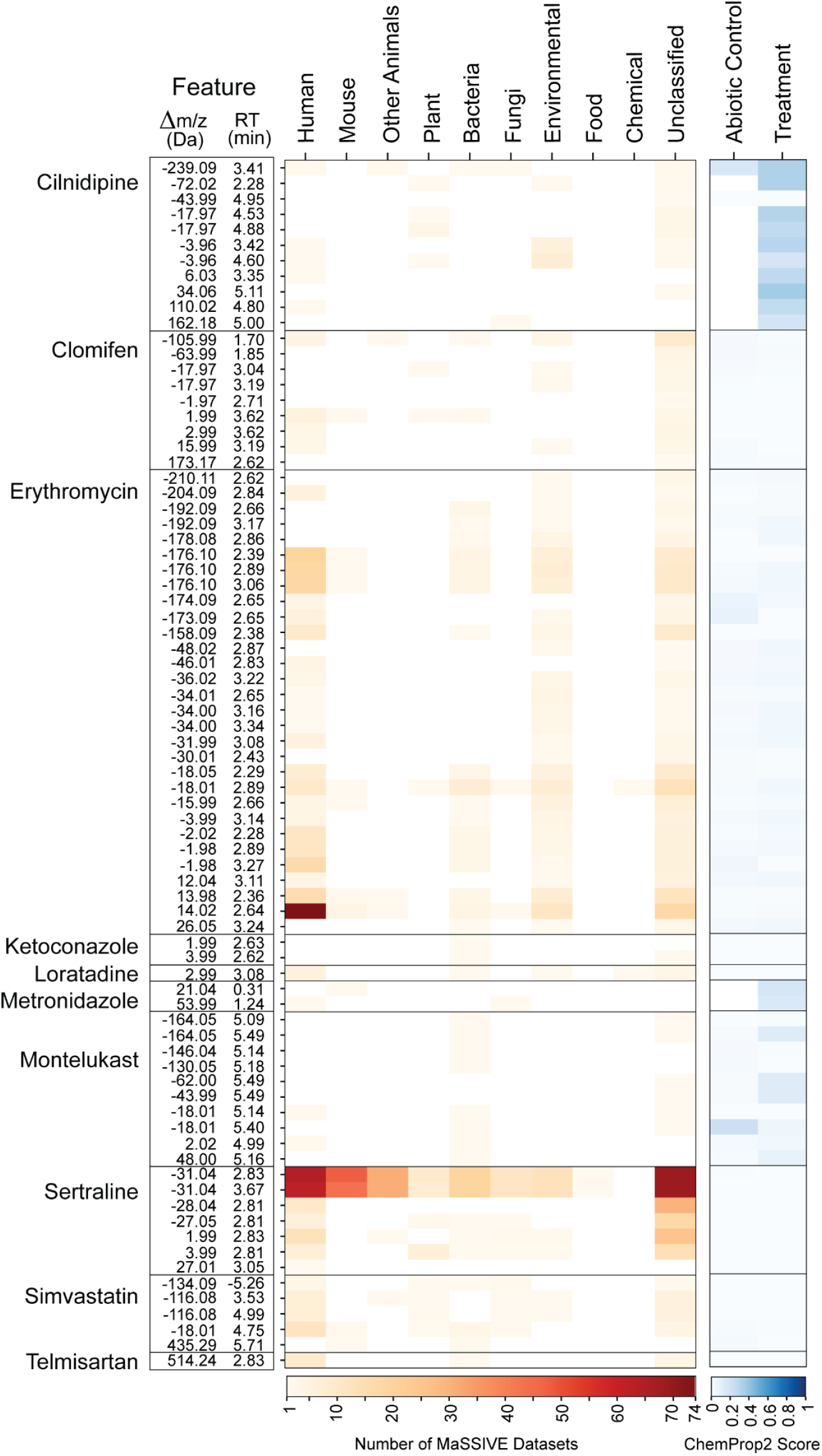
Prioritizing ChemProp2 Features Using Public Dataset Context. Heatmap showing the distribution of 78 ChemProp2-prioritized features (|ChemProp2 treatment score| > 0, |Δ*m/z*| > 0.5 Da, not unique to our dataset) across 10 high-level MassIVE dataset categories. Categories were derived from species-level metadata associated with matched GNPS entries. Corresponding ChemProp2 treatment scores for the same features, colored by score magnitude, illustrate directionality and potential transformation strength.

**Table 1:** ChemProp2 Summaries. The table provides the global subnetwork characteristics of the 12 drugs, including the number of cascade nodes, Δ*m/z* distributions, and annotation coverage from GNPS and FASST. ChemProp2 refined the interpretation of these otherwise unannotated nodes; for example, Sertraline yielded 39 prioritized features consistently detected across multiple datasets, while many Cilnidipine features were unique to this study yet exhibited strong treatment-associated trends (**Figure S14**).

| Drug Name | Nodes per drug | Nodes ($\Delta m/z > 0.5$) | Library Matched | Unmatched Compounds | MassIVE Matches | ChemProp2 Hits (>0.1) | Annotated Hits (>0.1) | MassIVE Hits (>0.1) | Final Hits |
| --- | --- | --- | --- | --- | --- | --- | --- | --- | --- |
| Cilnidipine | 98 | 98 | 0 | 98 | 20 | 76 | 0 | 14 | 11 |
| Clomifen | 42 | 36 | 1 | 37 | 34 | 9 | 0 | 6 | 9 |
| Erythromycin | 64 | 59 | 12 | 48 | 60 | 9 | 2 | 9 | 30 |
| Ketoconazole | 30 | 20 | 0 | 20 | 13 | 8 | 0 | 3 | 2 |
| Lansoprazole | 5 | 5 | 0 | 5 | 0 | 3 | 0 | 0 | 0 |
| Loratadine | 14 | 5 | 1 | 4 | 2 | 0 | 0 | 0 | 1 |
| Metronidazole | 18 | 6 | 0 | 6 | 3 | 3 | 0 | 2 | 2 |
| Montelukast | 70 | 64 | 0 | 64 | 27 | 10 | 0 | 5 | 10 |
| Omeprazole | 2 | 2 | 0 | 2 | 1 | 1 | 0 | 0 | 0 |
| Sertraline | 97 | 92 | 0 | 92 | 48 | 39 | 0 | 18 | 7 |
| Simvastatin | 55 | 53 | 18 | 38 | 38 | 4 | 2 | 7 | 5 |
| Telmisartan | 20 | 7 | 0 | 8 | 7 | 3 | 0 | 3 | 1 |
| **Total** | **515** | **447** | **32** | **422** | **253** | **165** | **4** | **67** | **78** |

Conversely, a +14.02 Da shift from Erythromycin matched across 74 human datasets and was annotated in GNPS as Clarithromycin, a broadly used clinical antibiotic derived from Erythromycin. In our time series, it showed a weak ChemProp2 score (0.02) and similar behavior in abiotic controls, suggesting a pre-existing analog rather than a microbially driven product.

Several strongly enriched features were not detected in any public repositories when queried (August 2025; **SI Figure 11**), suggesting that they may represent previously unreported metabolites unique to the biological conditions examined here. Together, this repository-level screening provides an important intermediate step between temporal inference and targeted experimental validation. By distinguishing recurrent metabolites from those unique to this system, public data help refine which candidate biotransformations warrant deeper mechanistic follow-up.

Contextualizing candidate biotransformations against public metabolomics repositories provides an important bridge between time-resolved inference and downstream biochemical validation. By examining whether features recur across independent datasets or remain exclusive to our system, we can prioritize metabolites that likely represent true microbiome-drug chemistry. For example, Cilnidipine produced 98 transformation nodes; 11 were also found in external datasets and exceeded the ChemProp2 treatment threshold, while 65 were unique to our experiment (60 of these also exceeded the threshold). This absence of public matches for several strongly enriched metabolites likely reflects the unique chemical space captured in our microbiome-drug system, as well as the natural variability in what has been deposited by the community to date. Additionally, upon prioritization with ChemProp2, complementary computational tools, such as ModiFinder^67^, can be used to help pinpoint the nature of the structure transformation, together providing a method combining MS/MS and time-series quantification. Combined with the appropriate computational tools to mine them, public repositories remain an invaluable resource, and as they continue to expand, some of these features may eventually find counterparts that help contextualize their biological origin.

## Discussion

This work demonstrates how *in vitro* experiments with the Com20 gut synthetic microbial community, combined with non-targeted metabolomics and our ChemProp data analysis approach, enable the systematic characterization of microbiome-mediated drug metabolism. Starting from 50 clinical drugs screened during endpoint experiments, we selected 12 compounds from different drug classes for a detailed timepoint analysis. Across these compounds, ChemProp revealed diverse, multi-step transformation patterns, including for Cilnidipine, Metronidazole, Omeprazole, and Simvastatin. Cascade expansion increased first-neighbor edges nearly six-fold, enabling detection of more distal products, which aligned with temporal changes in microbial composition.

Several metabolites appeared only at later timepoints and aligned with shifts in community composition; for example, late-stage Metronidazole derivatives coincided with a decrease of *Sarcina perfringens*, consistent with known previously described nitroreductase activity in gut-associated bacteria. These results demonstrate that microbial drug metabolism is not only shaped by the intrinsic chemistry of each drug but also by the ecological dynamics and functional capabilities of the resident microbes.

For other compounds, such as Cilnidipine, transformation patterns were consistent with both hepatic-like dehydrogenation and additional reductive processes attributed to gut bacterial activity, suggesting the coexistence of multiple metabolic routes. Similarly, the presence of multiple Omeprazole-derived products supports the involvement of anaerobic gut microbes in transformations extending beyond those typically associated with hepatic metabolism, with overlapping cascades potentially complicating directionality inference in complex communities.

To place these transformations in a broader biological context, repository-scale spectral searches helped distinguish metabolites observed in diverse hosts, from those exclusively observed in our *in vitro* experiments. This comparison highlights candidate metabolites potentially driven by the specific experimental system, while situating others within widely occurring populations. Beyond chemical diversification, microbial drug metabolism may influence both community structure and host drug exposure by altering compound activity, toxicity, or bioavailability.

Importantly, our chemical proportionality web application facilitates these analyses by enabling accessible computation of time-resolved ChemProp scores, cascade expansion, and interactive exploration of transformation networks. Integration with other computational tools in the GNPS2 environment, such as Modifinder, supports future structure-driven prioritization of candidate biotransformations. Together, these insights show how combining synthetic communities with time-resolved non-targeted metabolomics can disentangle microbial contributions to xenobiotic metabolism and reveal a more diverse, and dynamic chemical landscape than previously recognized.

## Online Methods

### Experimental designs and sample preparation

#### Endpoint screen

To investigate microbiome-mediated drug metabolism, 50 compounds were incubated with the Com20 synthetic gut community under anaerobic conditions in mGAM medium. Each drug was tested in triplicate at two timepoints (0 h and 2 h) alongside abiotic and vehicle controls. Incubations were performed in U-bottom 96-well plates (Thermo Fisher Scientific, cat. no. Z168136) and extracted with ethyl acetate prior to LC-MS/MS analysis. The screen was conducted in three batches based on drug solubility (aqueous, DMSO, or mixed).

#### Time-series experiment

Following the initial two-timepoint screening, a longitudinal experiment was conducted to assess temporal dynamics of microbial drug biotransformations and benchmark the ChemProp2 framework. Twelve compounds were selected: nine (Cilnidipine, Clomifen, Erythromycin, Ketoconazole, Lansoprazole, Loratadine, Metronidazole, Omeprazole, and Sertraline) that exhibited at least one ChemProp1 transformation edge above the threshold (score ≥ 1), and three (Montelukast, Simvastatin, and Telmisartan) included to test ChemProp2 performance in low-signal conditions.

Incubations were sampled hourly from 0 h to 8 h, each timepoint corresponding to an individual 96-well plate containing three biological replicates and one technical replicate per biological replicate. From each well, 900 μL was reserved for 16S rRNA sequencing. All drugs, except Cilnidipine and Loratadine, were tested at 8 μM; these two were used at 4 μM due to solubility constraints. Sample preparation and incubation conditions followed those used in the endpoint screening.

### Non-targeted Metabolomics using LC-MS/MS

Ethyl acetate extracts were dried under vacuum and resuspended in 50% methanol prior to LC-MS/MS analysis. Samples were measured on a Q Exactive HF Orbitrap mass spectrometer (Thermo Fisher Scientific) coupled to a Vanquish UHPLC system using a C18 column under a 5-minute reverse-phase gradient (0.1% formic acid in water/acetonitrile). Data were acquired in positive mode using data-dependent acquisition (DDA) with a resolution of 30,000 for MS^1^ and 15,000 for MS^2^. A pooled quality-control (QC) mix and an in-house six-compound QC mix were included to monitor retention time stability and batch effects. All chromatographic and MS parameters are provided in the Supplementary Methods.

### Data processing and FBMN

Raw LC-MS/MS data were converted to the .mzML format using ProteoWizard’s msconvert^83^, retaining only MS/MS scans. Files were processed in MZmine v4.0.3^84^ following a standardized batch workflow. Mass detection was performed separately for MS^1^ and MS^2^ scans using the Auto detector (noise level: 3 × 10^5^ and 1 × 10^3^, respectively). Chromatograms were built using the ADAP module (minimum five scans, height ≥ 1 × 10^6^, *m/z* tolerance = 0.002 Da or 10 ppm). Peaks were deconvoluted using the Minimum Search algorithm (duration 0.01-3 mins, minimum 5 data points, height ≥ 1 × 10^6^). MS^2^ spectra were linked to MS^1^ features within 10 ppm precursor *m/z* tolerance and retention time (RT) filtering. Isotopes were grouped using the Isotope Grouper and annotated via the Isotope Finder. Alignment across samples used the Join Aligner (10 ppm, 0.15 min RT tolerance), retaining features present in ≥ 3 samples and containing ≥ 2 isotopic peaks. Gap filling employed the Multithread Peak Finder (5 ppm, 0.05 min RT tolerance), and redundant features were removed using the Duplicate Filter (5 ppm, 0.1 min RT tolerance). Final feature tables containing MS^2^ scans (.mgf) and peak areas (.csv) were exported for GNPS(2) Feature-Based Molecular Networking (FBMN) and SIRIUS analysis.

FBMN was performed on GNPS(2) using default parameters unless stated otherwise. Precursor and fragment ion tolerances were both set to 0.01 Da; edges were retained for cosine > 0.7 with ≥ 6 matched peaks. Each node was connected to up to 10 most similar nodes (Δ*m/z* < 1999). Spectral library matching was performed against the GNPS library using identical thresholds, retaining only the top hit. A maximum component size of 100 was set. No intensity threshold or normalization was applied. The resulting edge table, together with the MZmine quantification table and sample metadata, was used as input for ChemProp analyses. GNPS job IDs, along with raw and processed data (.mzML), are listed in the Data Availability section.

### Com20 microbiome incubations and 16S rRNA sequencing

The *Com20* synthetic gut microbial community as describred in Griesshammer et al.^24^, comprises 20 commensal bacterial species spanning six phyla and representing ∼61% of the metabolic pathways found in a healthy human gut microbiome. All strains were originally obtained from DSMZ, BEI Resources, ATCC, or collaborating laboratories^24^ and routinely cultivated in pre-reduced mGAM medium (HyServe GmbH & Co. KG, Germany) at 37 °C under anaerobic conditions (2% H_2_, 12% CO_2_, 86% N_2_).

DNA from ChemProp2 time-series incubations (900 µL aliquots) was extracted using the DNeasy UltraClean 96 Microbial Kit (Qiagen). Amplicon sequencing of the 16S rRNA V4 region was performed on an Illumina MiSeq (2 × 250 bp) at the NGS Competence Center Tübingen following the protocol of Griesshammer et al^24^. Reads were processed with DADA2 (v1.21.0) and classified against a GTDB-based reference; Com20 members were further resolved to species level using a custom database (≥ 98 % identity). Species-level abundances were normalized by optical density (OD_600_) for downstream ordination (Bray-Curtis PCoA).

### ChemProp2 analysis and cascade scoring (including FDR)

ChemProp2 scores were computed using Pearson correlations across timepoints for each connected feature pair within the FBMN network. A strong anti-correlation between two features was interpreted as a potential precursor-product relationship, where one compound decreased and the other increased over time. To assess scoring reliability, ChemProp2 implements a false discovery rate (FDR)-based approach adapted from proteomics^46,47^ where peptide matches tested against real and decoy sequence databases are compared to estimate empirical thresholds. Feature tables are randomly shuffled to generate decoy datasets, which provide null distributions for comparison. Scores from original and decoy networks are then compared to estimate empirical FDR thresholds (e.g., 1%, 5%, 10%) for prioritizing high-confidence transformations. While this strategy improves interpretability, we note that shuffled datasets may still retain minor original patterns; further optimization of decoy generation will be addressed in future versions.

For cascade scoring, FBMN spectral library search returned 124 hits across the 12 analyzed drugs, including adducts and analogs. For downstream analysis, we focused on 14 features representing the parent [M+H]^+^ ions (**SI Table 6**). Most drugs had a single [M+H]^+^ feature, whereas Simvastatin (*m/z* 419.2791 and 419.2792; RT 4.75 and 5.01 min) and Telmisartan (*m/z* 515.2441 and 258.1256) contributed two each. The global FBMN with the 12 drugs contained 11,097 edges across 770 components, with 27 components including drug or analog nodes (2,155 edges total). Within this subset, 115 edges (D1) linked parent drug features to their immediate neighbors (**SI Table 3**). To capture downstream events, we expanded connections up to 10 cascade steps from each parent node. Cascade scoring added 3,637 edges, yielding 5,792 total scored edges (D1-D10). Of these, 664 edges were directly connected to 14 parent drug nodes, a ∼5.8-fold increase relative to D1 alone. This cascade expansion highlighted distal features with drug-linked temporal trends, offering a broader view of multi-step microbial biotransformation pathways. ChemProp2 scores were also computed for drug adduct and analogs and their neighbors, but the analyses presented in the manuscript were restricted to parent drug features.

### Selection of representative drugs for ChemProp2 analysis

From the initial drug screen, 50 compounds were selected for ChemProp1 analysis based on detectable precursor ions and network connectivity in FBMN. Each drug had a confidently observed [M+H]^+^ feature with adjacent nodes suitable for scoring, except for acarbose, detected primarily as an [M+NH4]^+^ adduct. ChemProp1 scores were calculated for every edge by comparing feature intensities between 0 h and 2 h, and edges with scores ≥ 1 (≥ two-fold change) were considered as candidate transformations.

Spectral library matches were found for 38 drugs (14 with class 1 annotations and 24 with class 3), providing additional confidence in compound identity. Parent ions were manually validated in the GNPS Dashboard by *m/z* and RT inspection. While some drugs displayed multiple network connections, the number of direct edges did not always correlate with transformation likelihood, highlighting the limitations of pairwise-only scoring. **Figure 2** summarizes these results, showing for each drug the number of first-degree connections and how many exceeded the ChemProp1 threshold. This initial screening established a baseline for assessing transformation potential and guided the selection of 12 representative drugs for subsequent time-resolved ChemProp2 analysis.

### Data Sharing

Raw LC-MS/MS data (.raw and .mzML) for all ChemProp1 experiments (50-drug screen, two timepoints) are publicly available on MassIVE (MSV000096724, MSV000093571) and Zenodo (https://zenodo.org/records/10213654, https://zenodo.org/records/10210429). The ChemProp2 multi-timepoint dataset (12 selected drugs) is available separately on MassIVE (MSV000094899) and Zenodo (https://zenodo.org/records/15677238). All molecular networking was conducted using the FBMN workflow on GNPS. Two GNPS1 jobs were used for the ChemProp1 screening experiment (https://gnps.ucsd.edu/ProteoSAFe/status.jsp?task=c561760343354873914a3f0bb4b03144, https://gnps.ucsd.edu/ProteoSAFe/status.jsp?task=421f28daa319440d900adfb2c9a56243). The longitudinal ChemProp2 dataset (12 drugs) was processed using the GNPS2 FBMN workflow (https://www.gnps2.org/status?task=1b5b94b4191d4223a5f57afb2aaaf0b0).

### Software Availability

The chemprop web application is publicly available for non-commercial use at https://chemprop.gnps2.org. A local installable application can be downloaded from https://www.functional-metabolomics.com/resources. All source code is publicly available at https://github.com/Functional-Metabolomics-Lab/ChemProp-Web-App and has been released as ChemProp2 v1.0.1.

### Use of Generative AI

No generative artificial intelligence (e.g. large language models) was used to generate original content of the manuscript. ChatGPT 5 (OpenAI) was used for proofreading and text editing of the manuscript. The authors take full responsibility for the content of the manuscript.

## Author Contributions

AKPS and DP conceptualized the ChemProp software. AKPS, AG, LM, and DP conceptualized the SynCom experiments. AKPS, AW, and MW developed the ChemProp application. AKPS, AG, and PS performed microbial culturing and extractions. PS performed mass spectrometry experiments. AG and LM performed amplicon sequencing. AKPS, AG, JCK, LM and DP analyzed data. AKPS and DP wrote the manuscript. All authors contributed to the writing and edited and approved the final manuscript.

## Acknowledgement

This study was supported by the Deutsche Forschungsgemeinschaft (German Research Foundation, DFG) via the Cluster of Excellence EXC 2124: Controlling Microbes to Fight Infection (CMFI, project ID 390838134) to LM and DP. PS was supported by the European Union’s Horizon Europe research and innovation programme through a Marie Skłodowska-Curie fellowship no. 101108450 MeStaLeM. JCK was supported by a Rhodes University postdoctoral fellowship. We further acknowledge support by the National Institute of General Medical Sciences, GM160154 to DP, and the National Institute of Diabetes and Digestive and Kidney Diseases, 5U24DK133658-02 to MW. LM acknowledges funding from the DFG (Emmy Noether Programme MA 8164/1-1), the ERC (gutMAP 101076967). We thank the NGS Competence Center Tübingen (NCCT).

## Competing interests

Mingxun Wang is a co-founder of Ometa Labs LLC.

