## Supplemental Information for "A Functional Metabolomics Framework to Track Microbiome Drug Metabolism"


### Supplementary Methods

#### Endpoint drug screening

To assess microbiome-mediated drug metabolism, 50 drugs were incubated with the Com20 synthetic gut community under anaerobic conditions in mGAM medium. Experiments were conducted in three batches based on drug solubility and replication design. All incubations were performed in U-bottom 96-well plates and extracted using ethyl acetate (EtOAc) at two timepoints (t = 0 h and 2 h). *Eggerthella lenta*, a slow-growing Com20 member, was pre-cultured several days in advance, while the remaining 19 strains were freshly inoculated in mGAM and incubated anaerobically at 37 °C. The Com20 community was assembled at an initial $OD_{578}$ of 0.01 in 5 mL mGAM and grown overnight at 37 °C under anaerobic conditions to reach OD 0.5-1.0.

Drug stocks were prepared in water or DMSO depending on solubility and diluted 1:20-1:50 in mGAM before plating. For DMSO-soluble drugs, 200 µL of diluted drug solution was dispensed into each well, followed by 200 µL of pre-grown Com20. For water-soluble drugs, 800 µL of Com20 was added directly to wells containing the diluted drug solution. Abiotic controls (drug only), vehicle controls (DMSO or water only), and at least two biological replicates per treatment were included on each plate.

Immediately after inoculation (t = 0 h), 400 µL from each well was extracted into 1 mL EtOAc, mixed, sealed, and stored at −20 °C. The remaining cultures were incubated anaerobically at 37 °C for 2 h, after which an additional 400 µL was extracted using the same procedure to generate t = 2 h samples.

### Non-targeted Metabolomics using LC-MS/MS

##### Sample extraction and preparation

Drug-microbiome incubations were performed under anaerobic conditions at the University Hospital Tübingen. At each timepoint, samples were collected in 96-deep-well plates (2 mL capacity), sealed, and stored at -80 °C until extraction. Frozen samples were transported on ice to the Functional Metabolomics Laboratory, University of Tübingen, for LC-MS/MS analysis and stored at -20 °C upon arrival. For extraction, plates were thawed at room temperature and mixed thoroughly before adding EtOAc. Each well contained 500 µL of sample, to which 1 mL of EtOAc was added. The mixtures were sonicated for 10 min and centrifuged at 3000 × g for 5 min. The upper EtOAc layer was transferred to new plates and dried under vacuum at room temperature using a SpeedVac concentrator. Dried extracts were resuspended in 150 µL of 50% methanol in water prior to LC-MS/MS measurement.

##### LC–MS/MS acquisition parameters

A pooled QC sample and an in-house six-compound QC mix were injected periodically to monitor instrument performance and retention time stability across runs. LC–MS/MS analyses were performed on a Q Exactive HF Orbitrap mass spectrometer (Thermo Fisher Scientific) equipped with a heated electrospray ionization (HESI) source and coupled to a Vanquish UHPLC system. Separation was achieved on a Kinetex C18 column (2.1 × 50 mm, 1.8 µm, 100 Å; Phenomenex). The mobile phases consisted of solvent A (water, LC/MS grade, Fisher Scientific) with 0.1% formic acid (FA), and solvent B (acetonitrile, LC/MS grade, Fisher Scientific) with 0.1% FA. After sample injection, a 5-minute linear gradient was applied. Solvent B was increased linearly from 5% to 50% over the first 4 minutes, then ramped to 99% between 4 and 5 minutes. This was followed by a 2-minute column wash out phase at 99% solvent B, and then a 2-minute column equilibration with 5% solvent B.

MS data were acquired in positive-ion mode using HESI (spray voltage 3.5 kV; sheath gas 50 AU; auxiliary gas 12 AU; sweep gas 1 AU; capillary 250 °C; aux heater 400 °C; S-lens RF level of 80 V). Full MS scans were collected from m/z 150-1500 *m/z* (resolution 30,000; AGC $1 \times 10^6$). Data-dependent MS/MS spectra were acquired for the top 5 intense precursor ions from each MS scan (resolution 15,000; AGC $5 \times 10^5$). An isolation window of 1 *m/z* was employed, followed by a stepped normalized collision energy (NCE) of 25, 35, 45 eV for ion fragmentation. Additional settings included a dynamic exclusion of 5.0 seconds, apex trigger enabled, and isotope exclusion enabled.

### ChemProp Webapp

The ChemProp web application supports CSV, XLSX, TSV, and TXT input formats and can also import data directly from GNPS2 via a Feature-Based Molecular Networking (FBMN) task ID. The interface includes modules for data preprocessing, ChemProp scoring, false discovery rate (FDR) analysis, and interactive visualization. Preprocessing options include blank removal, missing-value imputation, and total ion current (TIC) normalization.

ChemProp calculations can be performed in three complementary modes: (i) using a provided edge table, such as those derived from FBMN tasks or user-uploaded networks; (ii) through cascade analysis from a selected feature ID to explore higher-order transformation relationships; and (iii) by directly computing ChemProp scores between any two user-selected features. This flexibility allows users to evaluate transformations at different levels of network complexity, from targeted feature pairs to extended cascade structures.

ChemProp is available in two modes: ChemProp1, designed for endpoint (two-timepoint) datasets and producing unscaled scores, and ChemProp2, which supports multi-timepoint experiments and outputs normalized scores between 0 and 1. The resulting table reports node-pair information, including score magnitude (0 to 1) and transformation directionality (+1 or -1, indicating the predicted precursor-product relationship between connected features).

The FDR module, available only in ChemProp2, applies a target-decoy approach to estimate reliability thresholds (e.g., 1%, 5%, 10%). Randomized decoy feature tables are used to generate a null distribution for score comparison, and cumulative FDR values are computed across score bins. Users can then apply the FDR cutoffs to filter results, downloadable as a CSV file or as a Cytoscape-compatible ZIP archive containing GraphML and style files.

Visualization tools include the (i) global transformation plot, which displays ChemProp scores versus m/z differences to highlight transformation trends, and (ii) an interactive edge viewer that allows filtering by score range, m/z difference, annotation name, or node ID. Selected edges are visualized in two synchronized panels: (A) intensity profiles for the node pair across timepoints, and (B) the corresponding subnetwork with directional ChemProp arrows indicating putative precursor-product relationships. When input data are imported from an FBMN task ID, selected edges can additionally be inspected using GNPS Spectrum Resolver for spectral similarity assessment and Modifinder to explore plausible chemical modification sites.

#### Performance Comparison of ChemProp1 and ChemProp2

To compare ChemProp2 with ChemProp1, we reanalyzed the multi-timepoint dataset of 12 drugs using ChemProp1, restricting it to the end timepoints (T0 vs T8). ChemProp1 was applied only to direct drug neighbors (D1 edges) from the FBMN network, as cascade expansion was not part of its original framework. ChemProp1, which reports log fold-change between pairs of timepoints, yielded more non-zero edges since it captures any change over the entire period, regardless of fluctuations in between. However, ChemProp2 integrates all timepoints and applies FDR correction, resulting in fewer but more reliable drug-metabolite relationships. Overall, ChemProp1 detected 22 edges above the threshold (≥1), while ChemProp2 identified 36 edges above threshold 0.1 and 10 above threshold 0.3, corresponding to drug-specific FDR cutoffs (~±0.35-0.4). **Supplementary Table 3** summarizes, for each drug, the number of D1 edges retained by both methods at these thresholds, and **Supplementary Figure 13** illustrates this comparison for edges with Δm/z values between 0 and 200 Da.

#### FASST Searches

The FASST search was performed against 13 GNPS-indexed databases: NORMAN, ORNL_Bioscales2, ORNL_Populus_LC_MSMS, gnpsdata_index, gnpsdata_test_index, gnpslibrary, massivedata_index, massivekb_index, metabolomicspanrepo_index_latest, metabolomicspanrepo_index_nightly, panrepo_2024_11_12, panrepo_2025_07_09, panrepo_2025_08_06, and ptfi2_index (https://wang-bioinformatics-lab.github.io/GNPS2_Documentation/masst/, https://fasst.gnps2.org/libraries.

Across the FASST searches, matches were obtained from 1,670 unique MASSIVE datasets (MSV accessions). After excluding our six deposited repositories (MSV000093571, MSV000094899, MSV000096724, MSV000091452, MSV000095311, MSV000092059), hits remained in 1,664 external datasets.
Our FBMN analysis (GNPS2 job ID: 1b5b94b4191d4223a5f57afb2aaaf0b0), yielded 8,055 features, with 517 (6.4%) were annotated through GNPS spectral libraries, including all parent drug ions ($[M+H]^+$). ChemProp2 scores were calculated for all edges (cosine similarity-based and cascade). For downstream analysis, we focused on subnetworks centered around each drug's primary $[M+H]^+$ ion but also included treatment-specific features disconnected from the main clusters, as they may represent true biotransformation products. In total, 1,202 features were queried via FASST, comprising (i) nodes from $[M+H]^+$ clusters, (ii) adduct clusters, and (iii) treatment-specific features. FASST searches were performed with precursor tolerance 0.05 Da, fragment tolerance 0.05 Da, and cosine similarity ≥ 0.8. Both library hits and repository spectral matches were considered in downstream interpretation. Supplementary Table 4 summarizes FASST results by drug, indicating the number of features searched, those with at least one match, and the corresponding unique datasets (including and excluding our own repositories).

While all queried features were expected to return at least one match from our deposited datasets, 139 compounds did not. Manual verification through the FASST web interface (https://fasst.gnps2.org), confirmed that including precursor charge state restored matches for some cases. However, 120 features (16 from Cilnidipine, 104 from Simvastatin) still failed to return hits despite repeated manual checks. The cause remains unclear, and these 139 features were excluded from downstream analysis.

#### Cross-Dataset Distribution of Cascade Nodes (Heatmap Analysis)

As noted in the cascade summary, we began with 508 edges corresponding to drug-associated subnetworks and retained 433 cascade nodes with $\Delta m/z > 0.5$. To examine where these putative transformations are most prevalent, MassIVE metadata were retrieved for each feature and their species information were grouped into ten higher-level classes (e.g., Human, Mouse, Other Animals, Plant, Bacteria, Fungi, Environmental, Food, Chemical, Unclassified). Nodes were retained if they (i) had a non-zero ChemProp2 treatment score, (ii) differed from the parent drug by $\Delta m/z > 0.5$ Da, (iii) were observed in external MassIVE datasets, and (iv) represented forward transformations (Sign_ChemProp2_TRT = 1.0). This filtering resulted in 78 putative transformation nodes **(see Supplementary Table 5)**. The heatmap (**Figure 5**) shows their distribution across MassIVE dataset categories, alongside their ChemProp2 scores, highlighting the reproducibility and directional trend of the transformations.

### Cascade Node Summary Table

The following tables summarize drug-specific subnetworks, cascade edges, ChemProp2 hits, and whether features were annotated in GNPS libraries (FBMN) or found in external MassIVE repositories (FASST). Only cascade edges were included.

To generate these tables, we extracted all subnetworks from FBMN in which the parent drug $[M+H]^+$ ion resides (14 parent nodes in total: 12 drugs, with two $[M+H]^+$ nodes each for Simvastatin and Telmisartan, and one node for each of the remaining drugs). These 14 drug nodes all had library matches *(*See also the main manuscript section *Cascade Scoring Reveals Multi-Step Biotransformations*). For cascade scoring, we included only edges connecting each drug node to other nodes in the parent $[M+H]^+$ ion cluster, independent of spectral cosine similarity. In total, this yielded 515 drug-associated features. After removing the 14 parent drug ions, 436 features remained, connected by 508 edges with $|\Delta m/z| > 0.01$. Applying a more stringent filter of $|\Delta m/z| > 0.5$ reduced the set to 433 cascade nodes, which were used for the summary analyses below.

Each summary table contains the following fields:

- **Drug Name**: one of the 12 drugs.
- **Drug features**: number of parent drug nodes considered ($[M+H]^+$ only; 14 total).
- **Nodes (∆m/z > 0.5)**: number of cascade nodes differing from the drug node by more than 0.5 m/z. Nodes within ±0.5 m/z were excluded. For example, Cilnidipine yielded 98 such nodes.
- **Library Matched**: among these filtered nodes, the number of features with GNPS library matches.
- **Unmatched Compounds**: number of filtered nodes without library matches.
- **MassIVE Matches**: number of filtered nodes with ≥1 external hit in MassIVE datasets (counting nodes, not datasets).
- **ChemProp2 Hits (>0.1)**: number of filtered nodes with ChemProp2 treatment scores >0.1, indicating time-resolved intensity profiles consistent with downstream drug products.
- **Annotated Hits (>0.1)**: among ChemProp2 hits, number with GNPS library annotations.
- **MassIVE Hits (>0.1)**: among ChemProp2 hits, number also observed in external MassIVE datasets.
- **Final Hits:** Number of cascade nodes with ChemProp2 treatment score >0, m/z difference >0.5, significant treatment association (Sign_ChemProp2_TRT = 1.0) (This is represented as Heatmap in Fig 5 in the main manuscript)

### Supplementary Figures

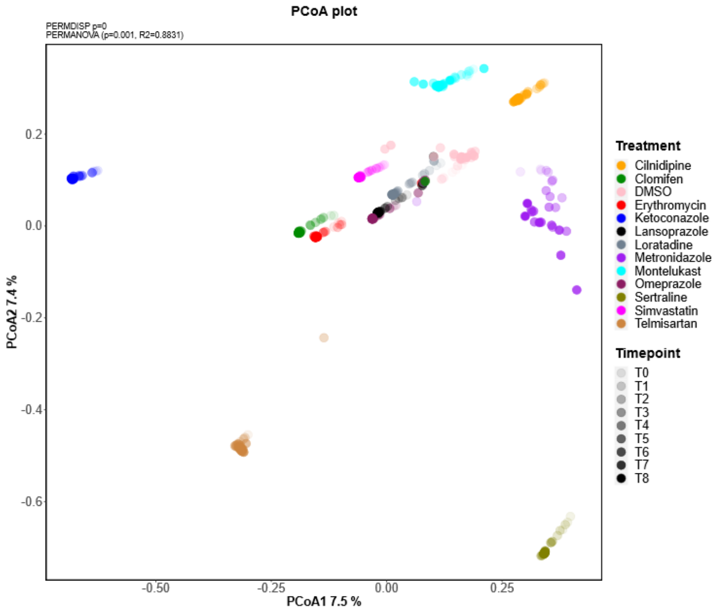


**Supplementary Fig. 1 |** PCoA (Bray-Curtis) of metabolomic profiles for the abiotic control. PERMANOVA and PERMDISP were performed across different drug treatments. PERMANOVA p = 0.001, $R^2$ = 0.88; PERMDISP p = 0.

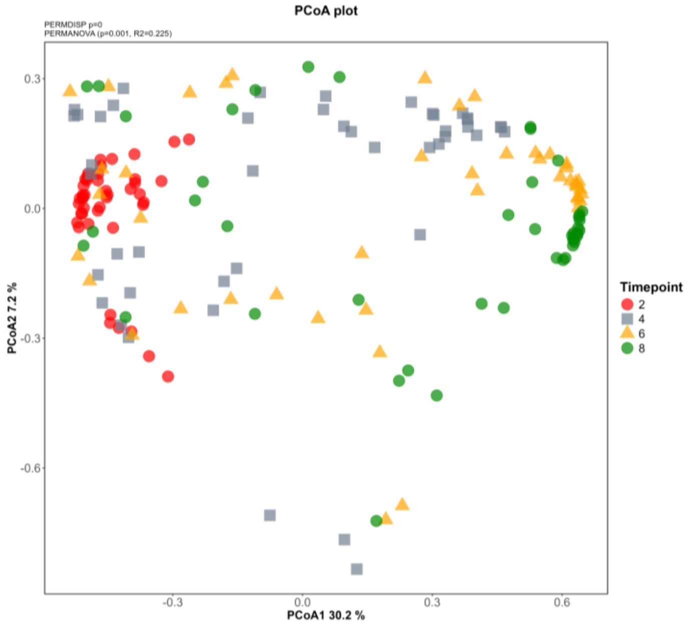


**Supplementary Fig. 2 |** PCoA (Bray-Curtis) of 16S rRNA ASV profiles for drug-treated bacterial communities. Points are colored by timepoint. PERMANOVA and PERMDISP were performed across T2, T4, T6, and T8 samples (PERMANOVA p = 0.001, $R^2$ = 0.225; PERMDISP p = 0).

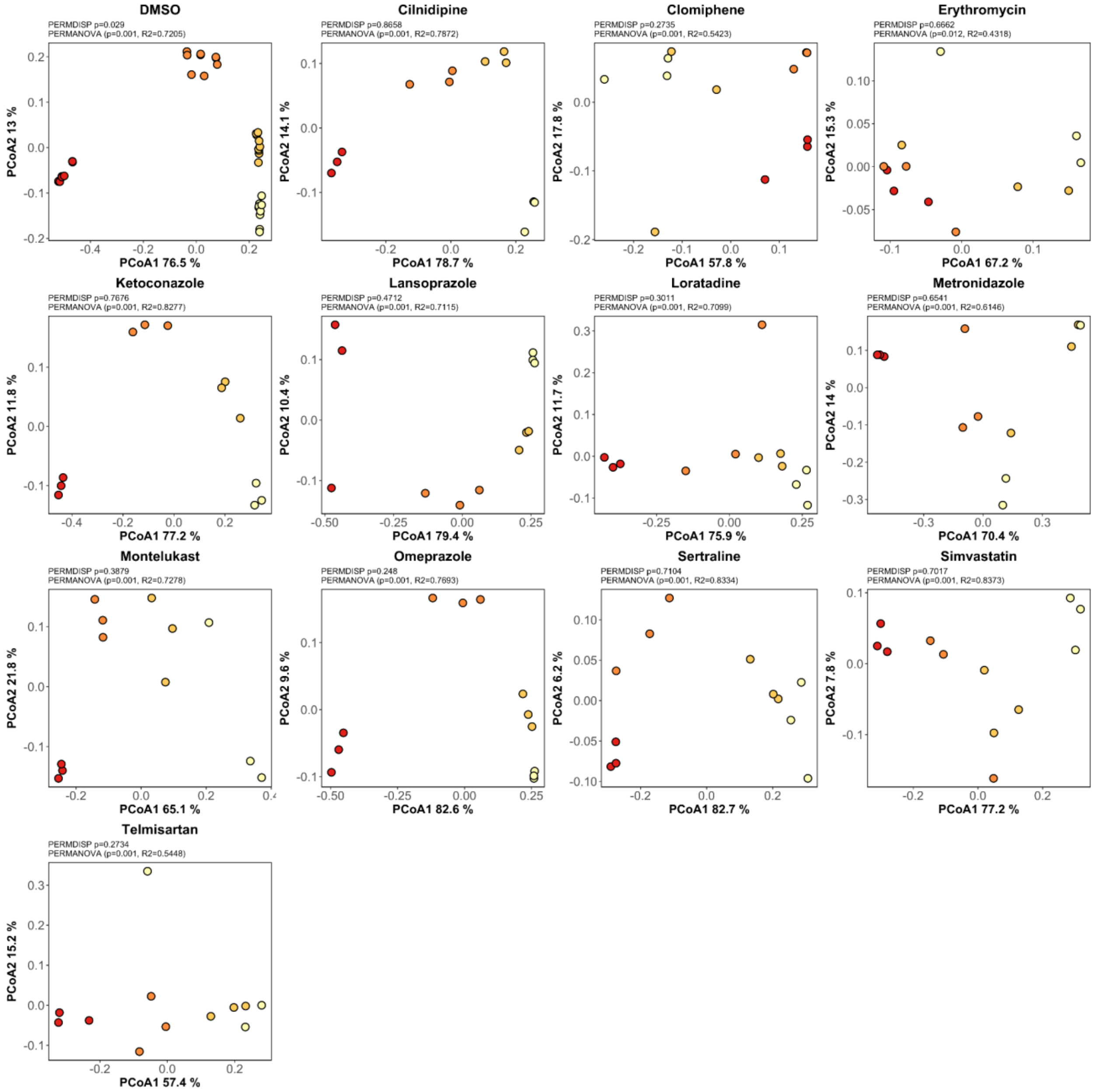


**Supplementary Fig. 3 |** Individual PCoA (Bray-Curtis) plots of OD-corrected microbiome profiles for Com20 treated with 12 different drugs and DMSO control (13 plots total). Points are colored from yellow (T2) to dark red (T8) to represent increasing timepoints. PERMANOVA and PERMDISP statistics shown in each plot are based on timepoint differences.

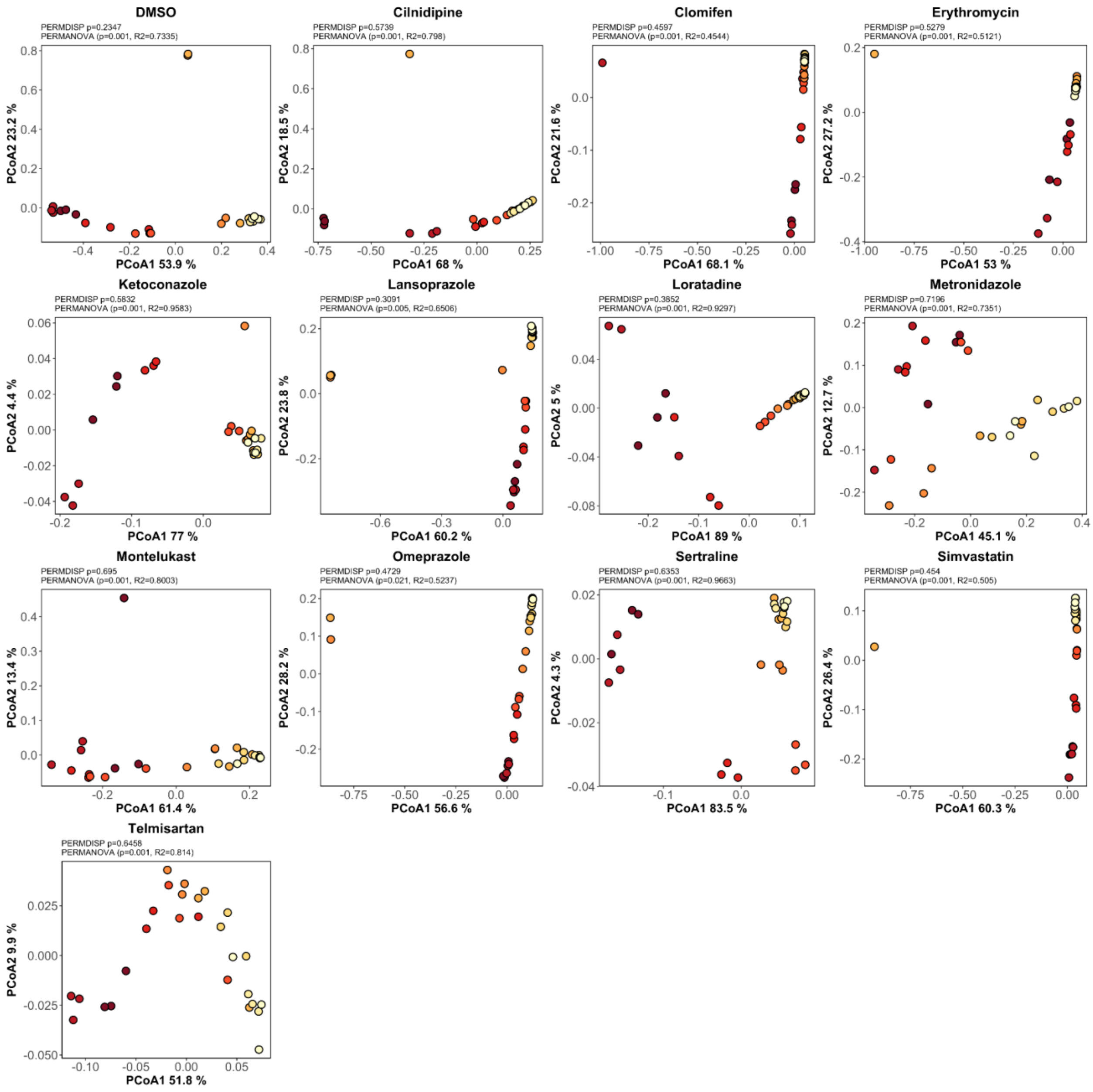


**Supplementary Fig. 4 |** Individual PCoA (Bray-Curtis) plots of metabolomic profiles for Com20 treated with 12 different drugs and DMSO control (13 plots total). Points are colored from light yellow (T0) to dark red (T8) to represent increasing timepoints. PERMANOVA and PERMDISP statistics shown in each plot are based on timepoint differences.

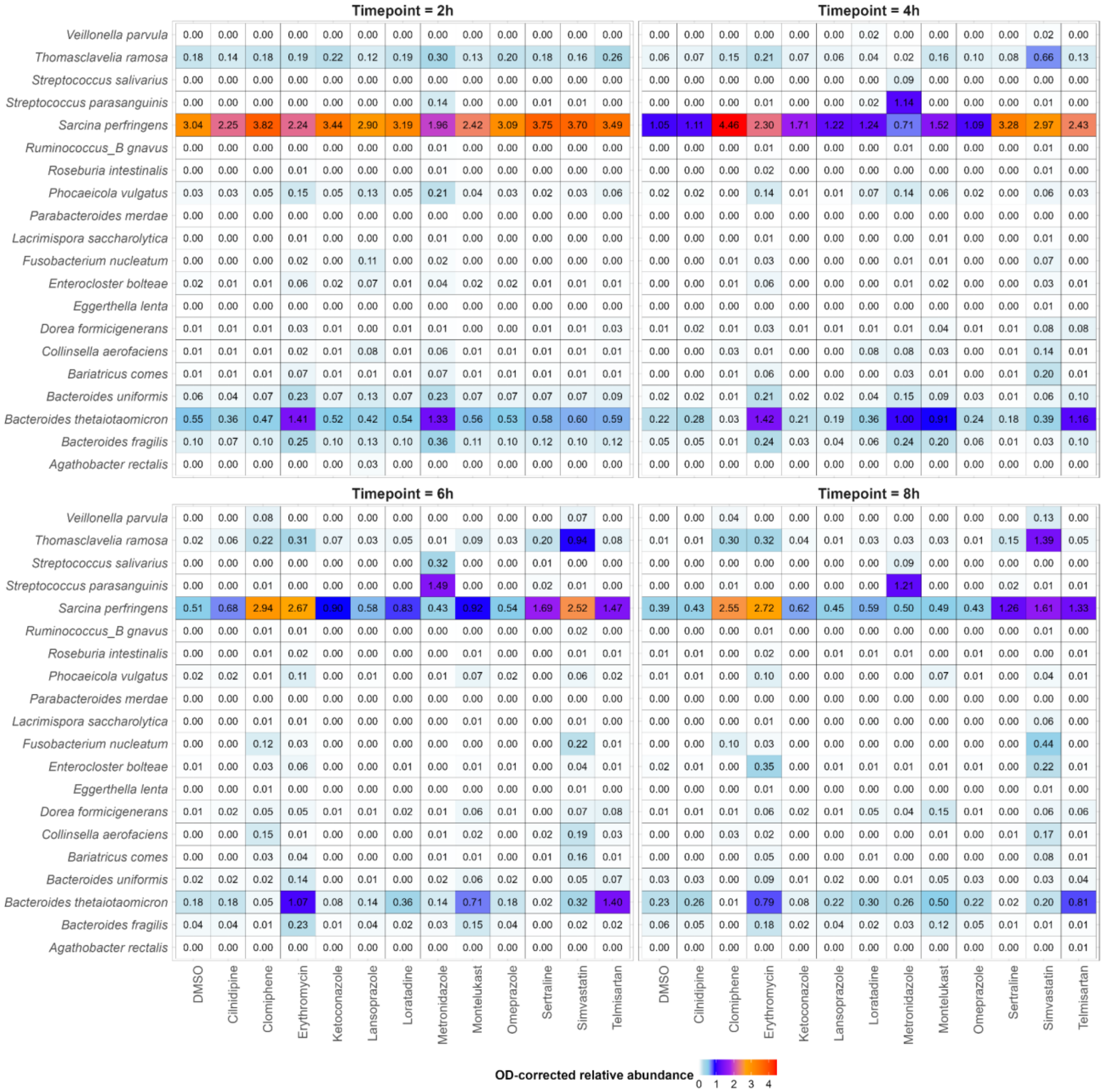


**Supplementary Fig. 5 | Heatmaps showing bacterial community responses across all drug treatments at each timepoint.** Mean OD-corrected abundances (averaged across replicates) are displayed for all 12 treatments. Rows correspond to individual taxa and columns to treatments. Color intensity reflects the OD-corrected mean relative abundance.

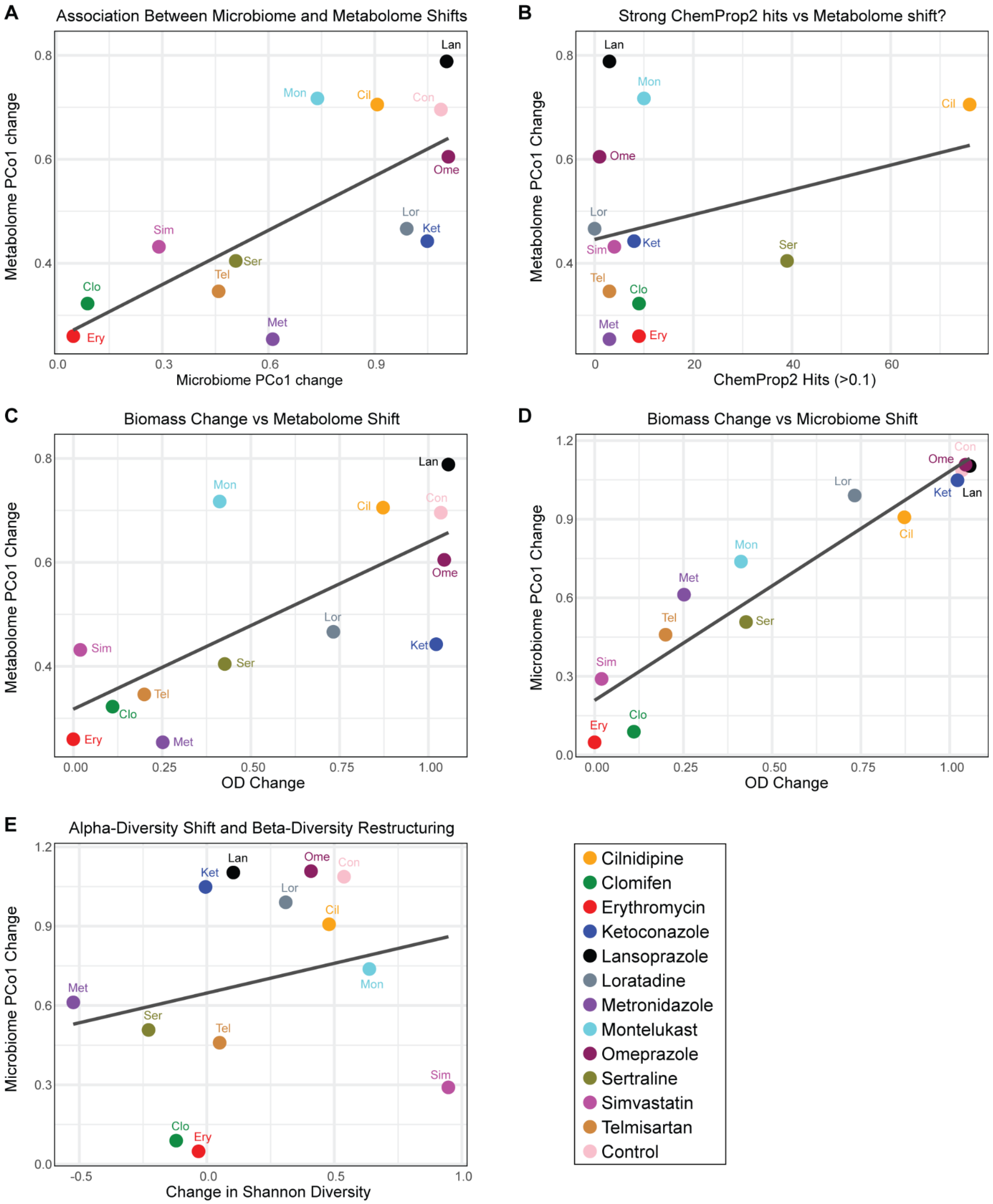


**Supplementary Fig. 6 | Multi-layer comparisons of drug-induced metabolome and microbiome shifts.** Each point represents one drug treatment, with metrics calculated from its time-series data (T0-T8). DMSO is included as the vehicle control and is summarized similar to drug treatments; all metrics were computed per treatment, and no comparisons or normalization were performed at individual timepoints relative to DMSO. **(A)** Relationship between metabolomic and microbiome restructuring over time. For each drug, PCo1 change reflects the total movement along the PCoA1 axis (metabolomics and 16S datasets), calculated as the cumulative change in mean PCoA1 values across consecutive timepoints; **(B)** Association between predicted chemical transformations and metabolomic change, showing the number of ChemProp2 hits (score > 0.1) per drug versus its PCoA1 trajectory length; **(C)** Biomass change versus metabolomic restructuring, comparing ΔOD (T8 – T2) with metabolome PCoA1 trajectory lengths for each drug; **(D)** Biomass change in relation to microbiome restructuring, with restructuring measured as compositional change captured by the microbiome PCoA1 trajectory length over time; **(E)** Diversity restructuring: alpha-diversity change (within-sample diversity from T0–T8) versus beta-diversity change (microbiome PCoA1 trajectory length).

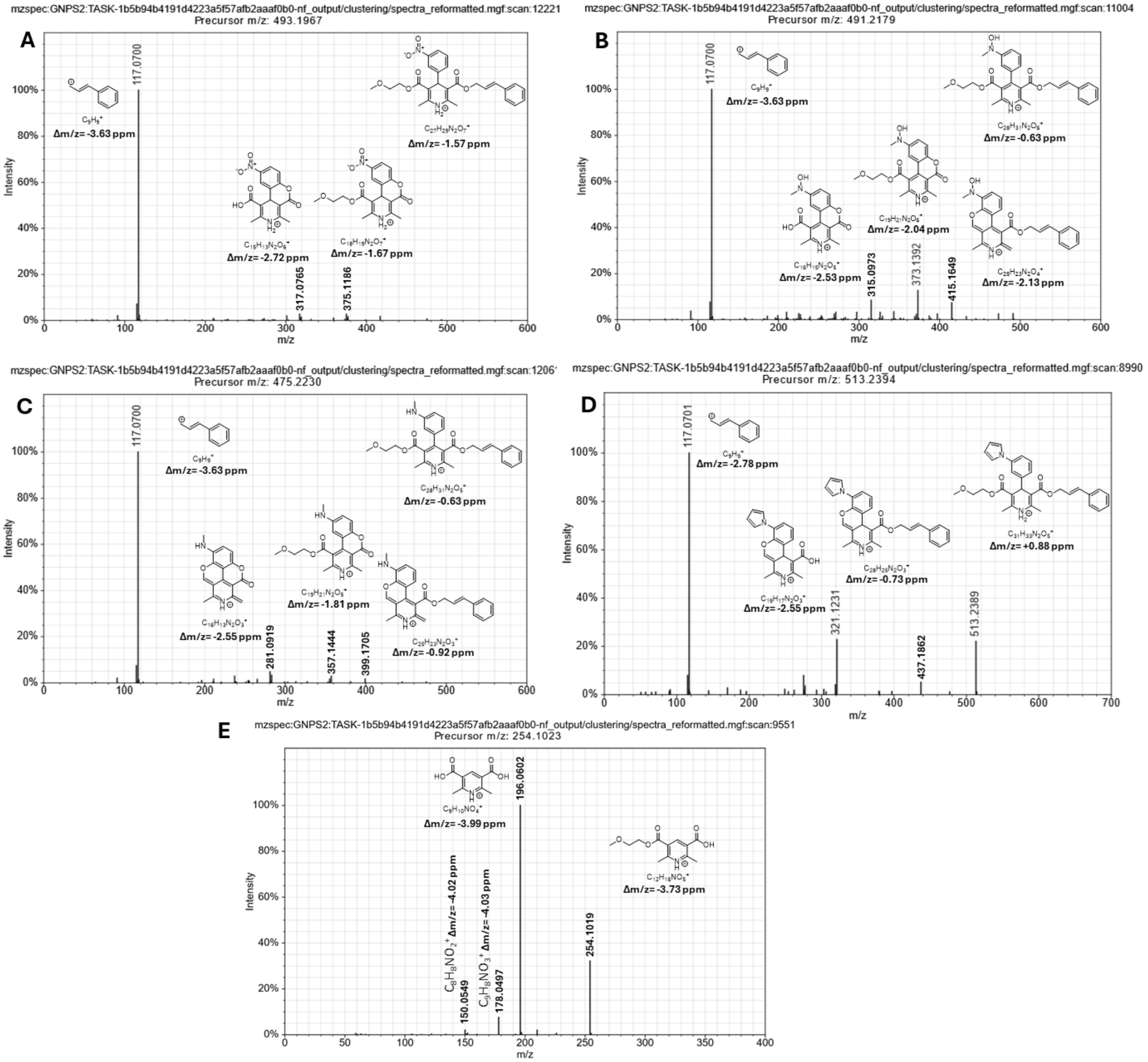


**Supplementary Fig. 7 | Fragmentation spectra and putative ion structure assignments for Cilnidipine and its putative microbial metabolites. A** Cilnidipine feature, **B** Putative Cilnidipine metabolite resulting from nitroreduction to a hydroxylamine and *N*-methylation, **C** Putative Cilnidipine metabolite resulting from complete nitroreduction an *N*-methylation, **D** Putative Cilnidipine metabolite resulting from nitroreduction and formation of a pyrrole moiety. **E** Putative Cilnidipine metabolite resulting from cleavage of the nitrobenzene moiety. Δ*m/z* values correspond to theoretical values subtracted by experimental values. Ion formulas were determined by accurate mass measurements and are unambiguous within the constraints of the assigned precursor molecular formulas. During CID the Cilnidipine related ions undergo fragmentation mechanisms involving rearrangement, though the exact nature of these remains unknown and the structures proposed by us only represent one set of possible structures as do the precursor structures for **B**, **C**, and **D**.

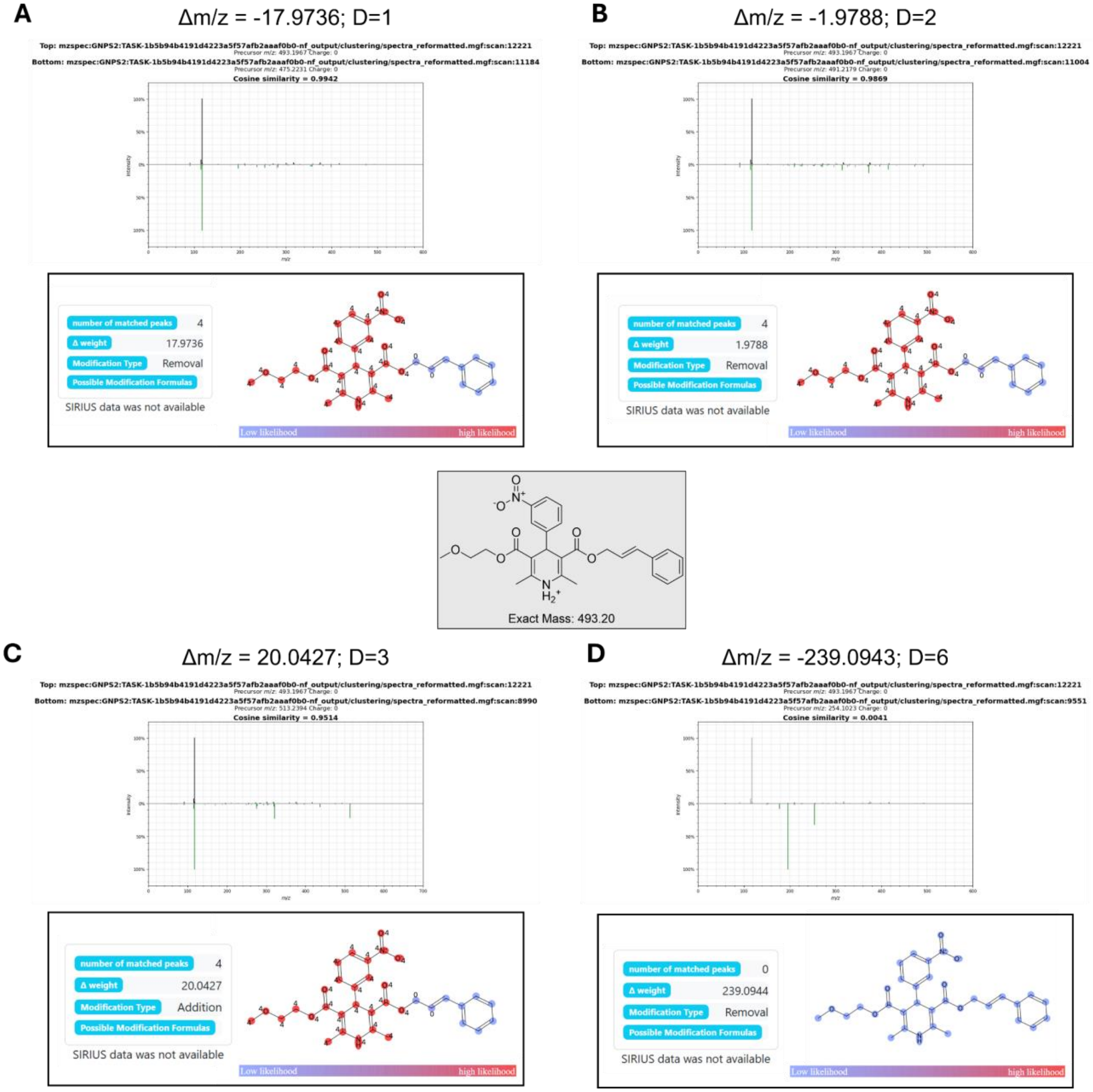

**Supplementary Fig. 8 | MS/MS spectral mirror matches and Modifinder predictions for ChemProp2-prioritized features of Cilnidipine.** The “D” values indicated in the panel titles along with m/z difference denote the degree of neighbor within each drug cluster.

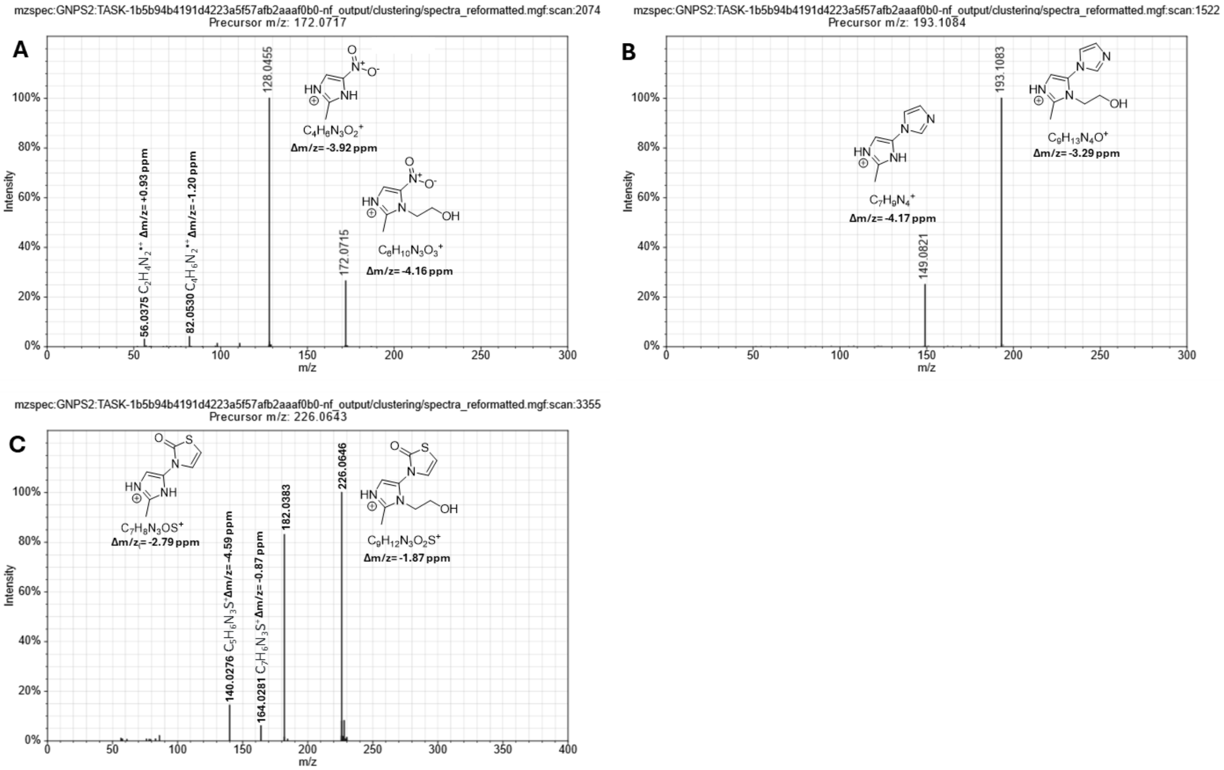


**Supplementary Fig. 9 | Fragmentation spectra and putative ion structure assignments for Metronidazole and its putative microbial metabolites. A** Metronidazole feature, **B** Putative Metronidazole metabolite resulting from nitroreduction and potential formation of an imidazole group **C** Putative Metronidazole metabolite resulting from nitroreduction and thiazolidinone formation, Δ*m/z* values correspond to theoretical values subtracted by experimental values. Ion formulas were determined by accurate mass measurements and are unambiguous within the constraints of the assigned precursor molecular formulas. Molecular structures for fragment ions and potential microbial metabolite precursors are putative and cannot be confidently ascertained based on the current data.

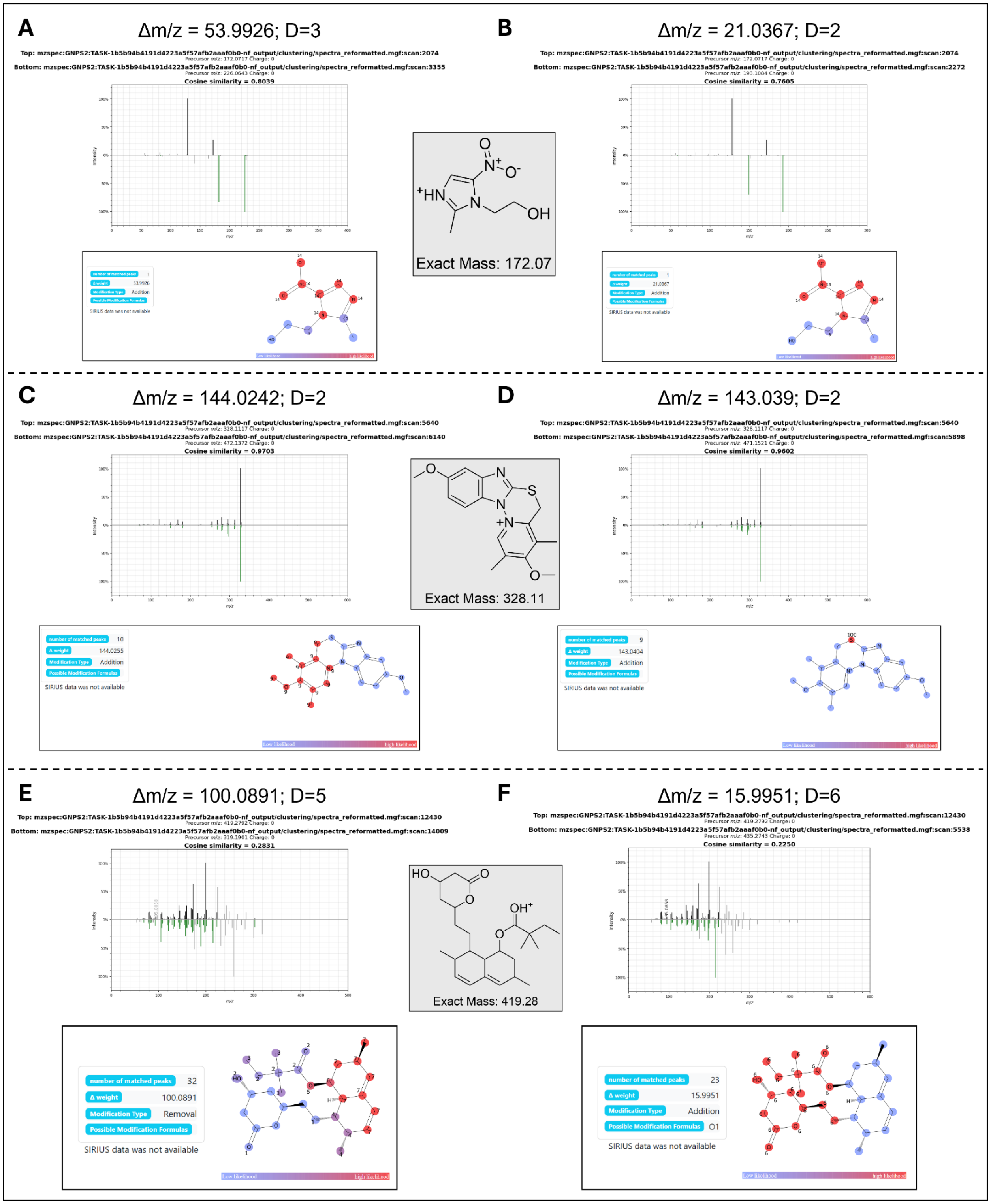


**Supplementary Fig. 10 | MS/MS spectral mirror matches and Modifinder predictions for ChemProp2-prioritized features.** Panels **A-B** show metronidazole features; **C-D** show omeprazole (M+H-H2O); and **E-F** show simvastatin (M+H). The "D" values indicated in the panel titles denote the degree of neighbor within each drug cluster.

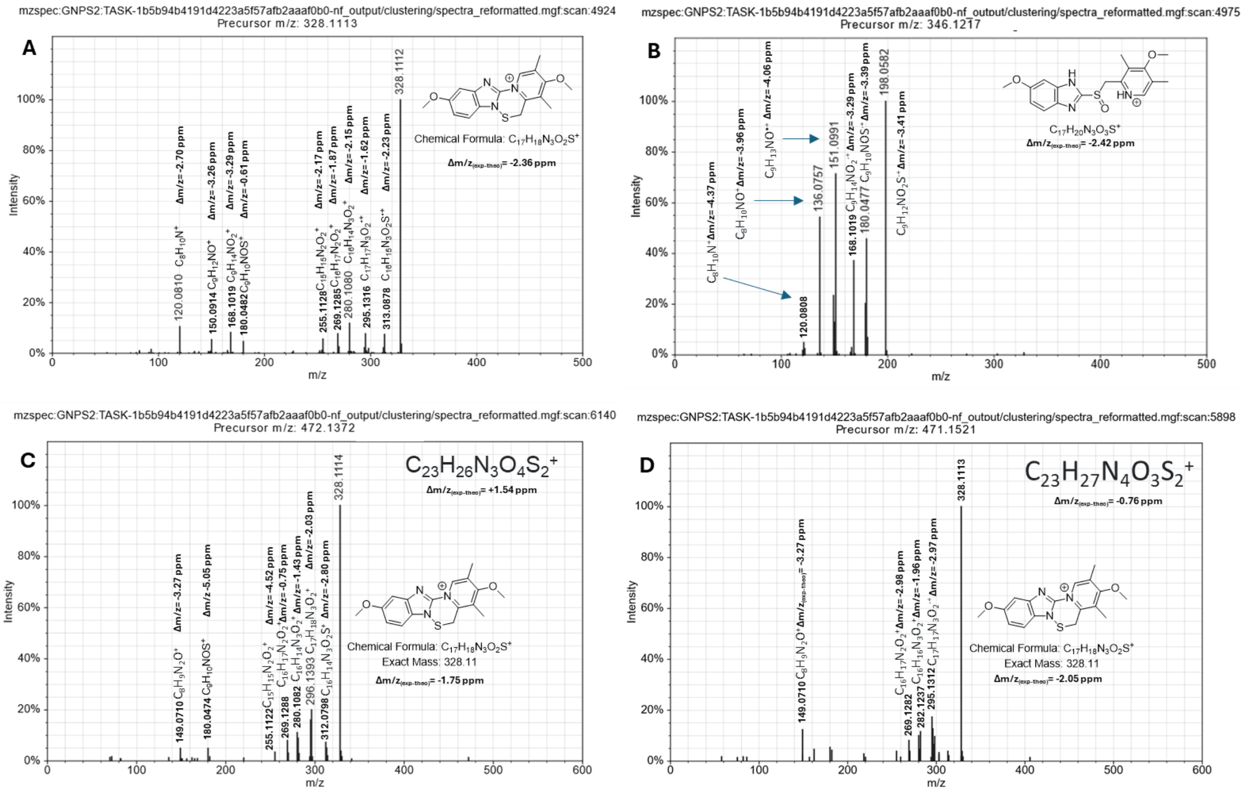


**Supplementary Fig. 11 | Fragmentation spectra and putative ion structure assignments for Omeprazole and its putative in-source fragment and microbial metabolites. A** Omeprazole in-source fragment feature, **B** Omeprazole feature, **C** Putative Omeprazole metabolite resulting from addition of $C_6H_6OS$, **D** Putative Omeprazole metabolite resulting from addition of $C_6H_7NS$. Δ*m/z* values correspond to theoretical values subtracted by experimental values. Ion formulas were determined by accurate mass measurements and are unambiguous within the constraints of the assigned precursor molecular formulas. Putative Omeprazole metabolites produce major fragment ions at *m/z* 328, matching the omeprazole in-source fragment and indicating neutral loss of all elements added during hypothetical biotransformation. During CID the Omeprazole related ions appear to undergo fragmentation mechanisms involving rearrangement, complicating evidence-based assignment of fragment structures.

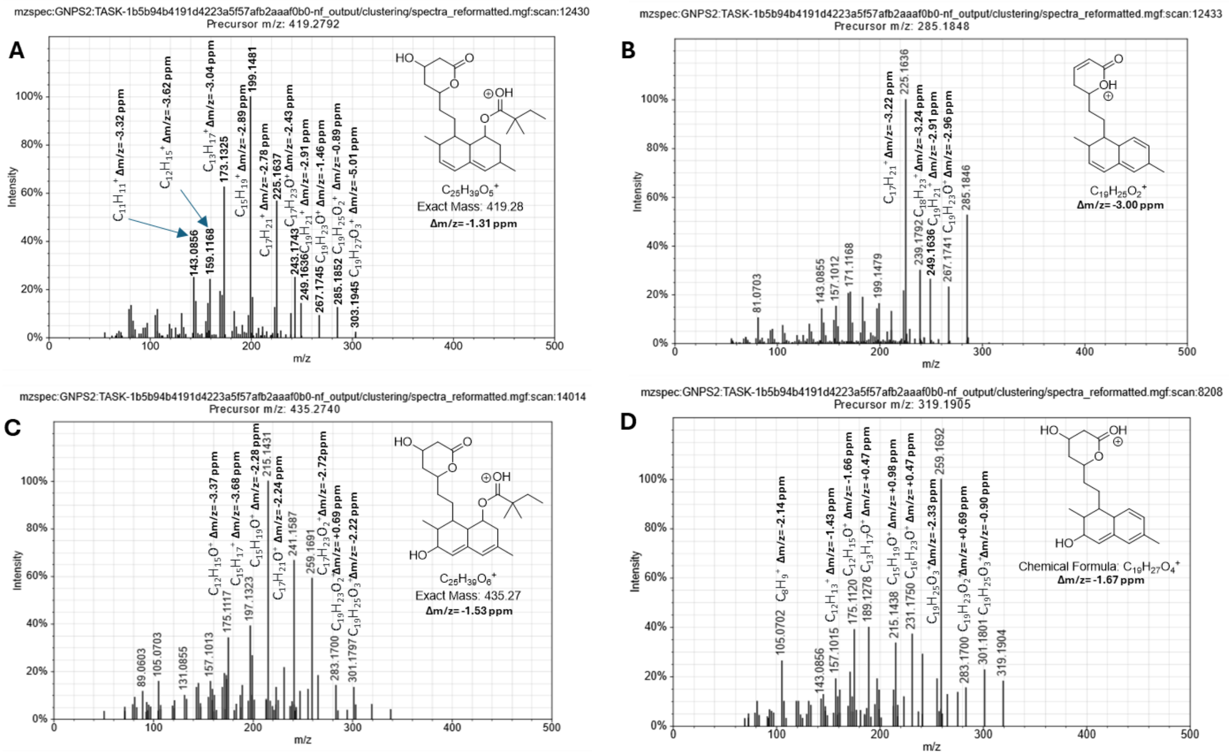


**Supplementary Fig. 12 | Fragmentation spectra and putative ion structure assignments for Simvastatin, its putative microbial metabolite and their in-source fragments. A** Simvastatin feature, **B** Simvastatin in-source fragment feature, **C** Putative Simvastatin metabolite 3'-Hydroxysimvastatin **D** Putative in-source fragment of 3'-Hydroxysimvastatin. Δ*m/z* values correspond to theoretical values subtracted by experimental values. Ion formulas were determined by accurate mass measurements and are unambiguous within the constraints of the assigned precursor molecular formulas.

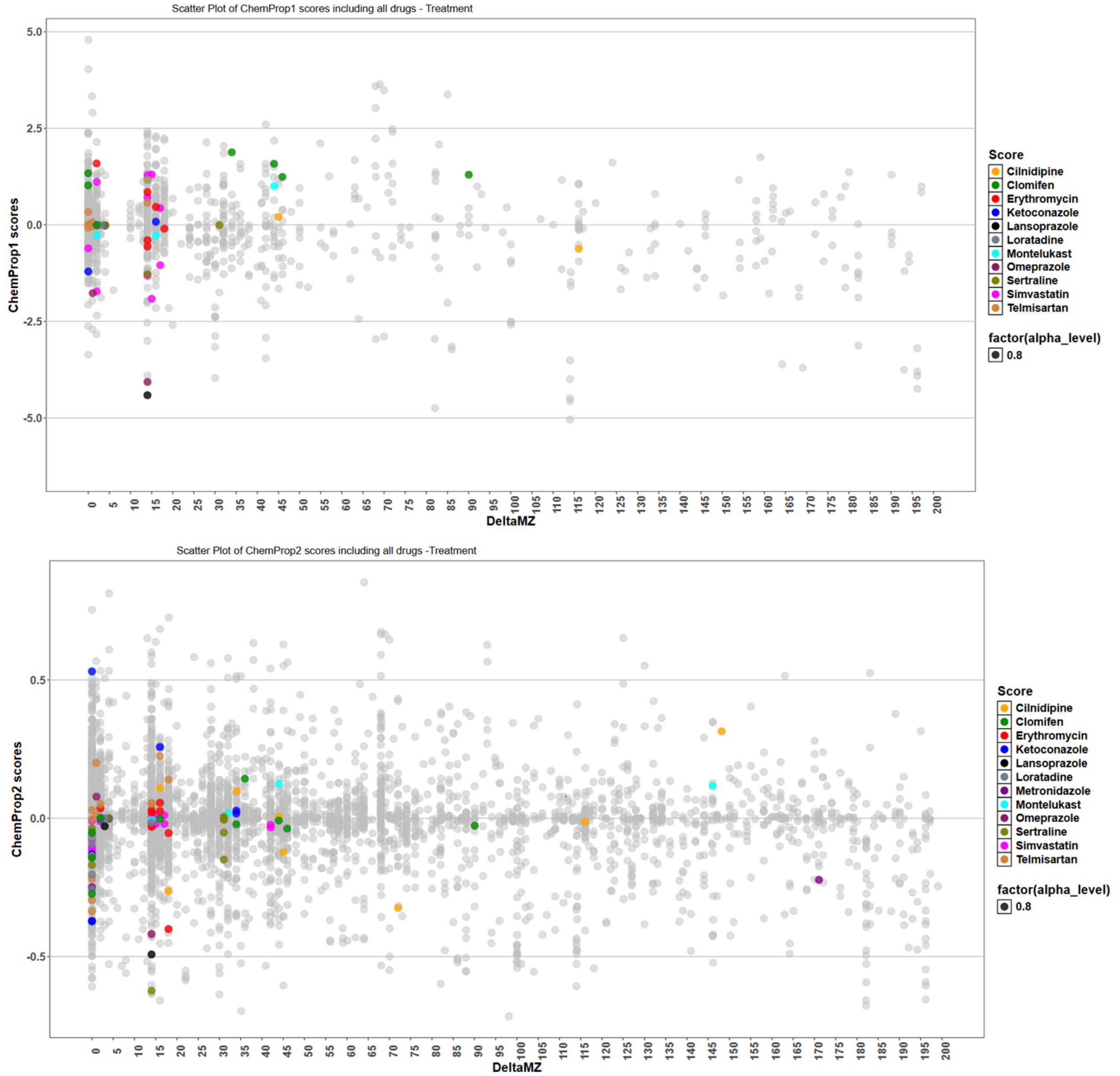


**Supplementary Fig. 13 | Scatterplot of ChemProp Scores for 12 Drugs.** The plot shows ChemProp scores (y-axis) against Δm/z values (x-axis) for first-degree (D1) edges connected to each drug, with points colored by drug as indicated in the legend. The comparison between ChemProp1 and ChemProp2 was restricted to edges with Δ*m/z* values between 0 and 200 Da.

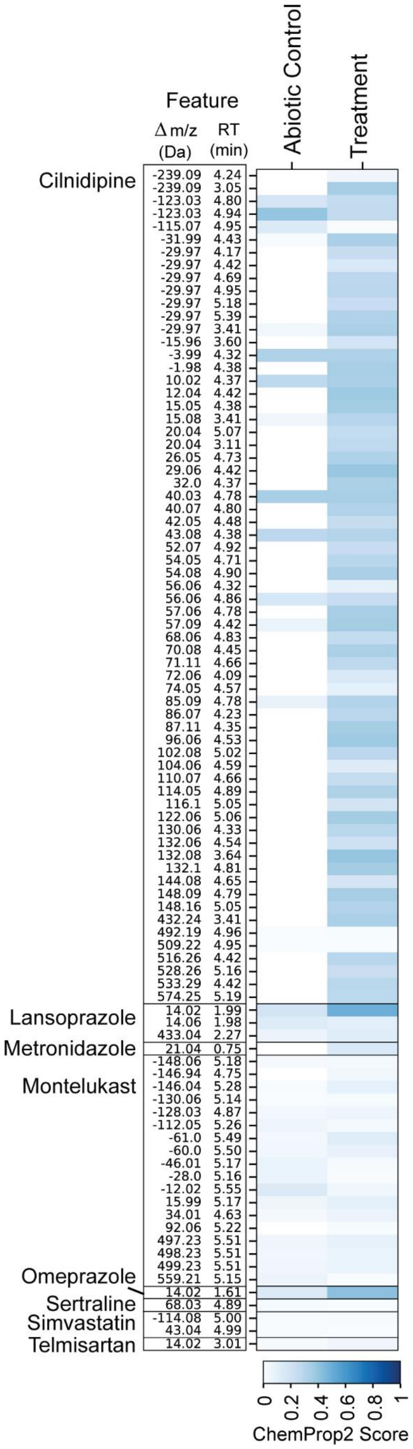


**Supplementary Fig. 14 |** ChemProp2-prioritized features unique to our dataset

### Supplementary Tables

**Supplementary Table 1 |** Drug class abbreviations for the 50 compounds analyzed using ChemProp1 (corresponding to Figure 2). Classes are grouped into six major categories, and cell colors correspond to the drug classes shown in **Figure 2**.

| Drug Class | Code | Abbreviation |
|---|---|---|
| Antibiotics & antiparasitics | AB | Antibiotic |
| | ANTIDIAB | Antidiabetic |
| | APAR | Antiparasitic |
| Antifungals | AF | Antifungal |
| Anti-inflammatory/Others | HORM | Hormonal |
| | NSAID | Nonsteroidal Anti-inflammatory Drugs |
| | OTHER | Other / Miscellaneous Drugs |
| | PPI | Proton Pump Inhibitors |
| | SERM | Selective Estrogen Receptor Modulators |
| Cardiovascular | ANTICO | Anticoagulants |
| | ARB | Angiotensin II Receptor Blockers |
| | CCB | calcium-channel blocker |
| | STATIN | statins |
| CNS/ Psychotrophic | AD | Antidepressant |
| | AP | Antipsychotic |
| | NK1 | Neurokinin-1 Receptor Antagonists |
| | SNRI | Serotonin-Norepinephrine Reuptake Inhibitors |
| | SSRI | Selective Serotonin Reuptake Inhibitors |
| Allergy/Immune | AH | Antihistamines |
| | LTRA | Leukotriene Receptor Antagonists |

**Supplementary Table 2 |** Drug-specific summary of cascade composition, microbial diversity, biomass changes, and Bray-Curtis PCoA shifts.

| Drug Name | Nodes per drug | Nodes (m/z > 0.5) | Library Matched | Unmatched Compounds | MassIVE Matches | ChemProp2 Hits (>0.1) | Final Hits | Microbiome α (mean) | Microbiome α (SD) | Microbiome change in α | ΔOD | OD mean | OD SD | Microbiome change in β (PCo1) | Microbiome change in β (PCo2) | Metabolome change in β (PCo1) | Metabolome change in β (PCo2) |
|---|---|---|---|---|---|---|---|---|---|---|---|---|---|---|---|---|---|
| Cilnidipine | 98 | 98 | 0 | 98 | 20 | 76 | 11 | 1.12 | 0.20 | 0.48 | 0.87 | 0.80 | 0.35 | 0.91 | 0.28 | 0.71 | 0.11 |
| Clomifen | 42 | 36 | 1 | 37 | 34 | 9 | 9 | 0.64 | 0.39 | -0.12 | 0.11 | 0.25 | 0.06 | 0.09 | 0.21 | 0.32 | 0.19 |
| Erythromycin | 64 | 59 | 12 | 48 | 60 | 9 | 30 | 1.46 | 0.15 | -0.03 | 0.00 | 0.21 | 0.01 | 0.05 | 0.14 | 0.26 | 0.15 |
| Ketoconazole | 30 | 20 | 0 | 20 | 13 | 8 | 2 | 0.79 | 0.15 | 0.00 | 1.02 | 0.72 | 0.41 | 1.05 | 0.32 | 0.44 | 0.11 |
| Lansoprazole | 5 | 5 | 0 | 5 | 0 | 3 | 0 | 1.07 | 0.17 | 0.10 | 1.06 | 0.86 | 0.45 | 1.10 | 0.47 | 0.79 | 0.13 |
| Loratadine | 14 | 5 | 1 | 4 | 2 | 0 | 1 | 1.07 | 0.23 | 0.31 | 0.73 | 0.63 | 0.29 | 0.99 | 0.12 | 0.47 | 0.09 |
| Metronidazole | 18 | 6 | 0 | 6 | 3 | 3 | 2 | 1.46 | 0.41 | -0.52 | 0.25 | 0.34 | 0.11 | 0.61 | 0.70 | 0.25 | 0.11 |
| Montelukast | 70 | 64 | 0 | 64 | 27 | 10 | 10 | 1.39 | 0.26 | 0.64 | 0.41 | 0.45 | 0.19 | 0.74 | 0.30 | 0.72 | 0.11 |
| Omeprazole | 2 | 2 | 0 | 2 | 1 | 1 | 0 | 1.12 | 0.17 | 0.41 | 1.04 | 0.85 | 0.44 | 1.11 | 0.34 | 0.60 | 0.10 |
| Sertraline | 97 | 92 | 0 | 92 | 48 | 39 | 7 | 0.61 | 0.17 | -0.23 | 0.43 | 0.40 | 0.18 | 0.51 | 0.28 | 0.40 | 0.09 |
| Simvastatin | 55 | 38 | 18 | 55 | 53 | 4 | 5 | 1.38 | 0.41 | 0.95 | 0.02 | 0.21 | 0.01 | 0.29 | 0.03 | 0.43 | 0.15 |
| Telmisartan | 20 | 7 | 0 | 8 | 7 | 3 | 1 | 1.07 | 0.15 | 0.05 | 0.20 | 0.30 | 0.09 | 0.46 | 0.36 | 0.35 | 0.10 |
| Control | | | | | | | | 1.13 | 0.23 | 0.54 | 1.04 | 0.92 | 0.43 | 1.09 | 0.41 | 0.70 | 0.18 |

**Supplementary Table 3 | Comparison of ChemProp1 and ChemProp2 D1 Edges Across Thresholds for Each Drug.**
D1 represents direct connections (edges) to the parent drug node in the FBMN network, whereas all other connections beyond the first-degree neighbor are referred to as global edges. Cell values are color-coded from red to white to blue, indicating high, medium, and low values, respectively.

| Drug | D=1 Edges | CHEMPROP 1 | CHEMPROP2 | | |
|---|---|---|---|---|---|
| | | Edges >= 1 | Edges != 0 | Edges > ±0.1 | Edges > ±0.3 |
| Cilnidipine | 10 | **0** | 10 | **5** | 2 |
| Clomifen | 10 | **6** | 10 | **3** | 0 |
| Erythromycin | 8 | **1** | 8 | **1** | 1 |
| Ketoconazole | 10 | **1** | 9 | **5** | 3 |
| Lansoprazole | 4 | **1** | 3 | **2** | 1 |
| Loratadine | 10 | **0** | 8 | **3** | 0 |
| Metronidazole | 6 | **0** | 2 | **2** | 0 |
| Montelukast | 10 | **2** | 10 | **2** | 0 |
| Omeprazole | 2 | **2** | 2 | **1** | 1 |
| Sertraline | 10 | **1** | 10 | **3** | 1 |
| Simvastatin | 19 | **7** | 19 | **2** | 0 |
| Telmisartan | 16 | **1** | 15 | **7** | 1 |
| Global | 10982 | **287** | 5922 | **1833** | 519 |

| | | | | | |
|---|---|---|---|---|---|
| Total | 11097 | **309** | 6028 | **1869** | 529 |

**Supplementary Table 4 | Summary of FASST results for ChemProp2-prioritized features.** Columns show the number of features queried per drug, features with at least one match, and those without matches. Also reported are the counts of unique datasets matched across all repositories, as well as subsets limited to MassIVE and external MassIVE datasets (excluding our six deposited repositories). “All Unique Datasets Matched” values are provided per drug but not summed in the summary row.

| **Drug** | **Features FASST input** | **Features with ≥ 1 Hit** | **Features without Hits** | **All Unique datasets Matched** | **Unique MASSIVE datasets** | **Unique External MassIVE Datasets** |
|---|---|---|---|---|---|---|
| Cilnidipine | 309 | 289 | 20 | 363 | 332 | 327 |
| Clomifen | 47 | 47 | 0 | 194 | 175 | 170 |
| Erythromycin | 113 | 113 | 0 | 221 | 119 | 194 |
| Ketoconazole | 39 | 37 | 2 | 52 | 50 | 47 |
| Lansoprazole | 102 | 101 | 1 | 290 | 257 | 251 |
| Loratadine | 16 | 16 | 0 | 75 | 66 | 63 |
| Metronidazole | 27 | 24 | 3 | 1144 | 1014 | 1010 |
| Montelukast | 83 | 78 | 5 | 456 | 400 | 395 |
| Omeprazole | 101 | 100 | 1 | 491 | 442 | 436 |
| Sertraline | 98 | 98 | 0 | 487 | 413 | 409 |
| Simvastatin | 245 | 141 | 104 | 829 | 772 | 766 |
| Telmisartan | 22 | 19 | 3 | 68 | 57 | 54 |
| **Total** | 1202 | 1063 | 139 | not calculated | 1670 | 1664 |

**Supplementary Table 5 | Summary of cascade node counts after ChemProp2 filtering.** Counts of external and internal nodes per drug, with corresponding ChemProp2 treatment scores. External refers to features detected in external datasets, whereas Internal nodes were found only in our six in-house datasets.

| **Drug** | **Cascade Nodes per drug** | **Nodes (Δm/z > 0.5)** | **Nodes After ChemProp2 Filtering (External)** | **Nodes After ChemProp2 Filtering (Internal)** |
|---|---|---|---|---|
| Cilnidipine | 98 | 98 | 11 | 65 |
| Clomifen | 42 | 36 | 9 | 0 |
| Erythromycin | 64 | 59 | 30 | 0 |
| Ketoconazole | 30 | 20 | 2 | 0 |
| Lansoprazole | 5 | 5 | 0 | 3 |
| Loratadine | 14 | 5 | 1 | 0 |

| | | | | |
|---|---|---|---|---|
| Metronidazole | 18 | 6 | 2 | 1 |
| Montelukast | 70 | 64 | 10 | 18 |
| Omeprazole | 2 | 2 | 0 | 1 |
| Sertraline | 97 | 92 | 7 | 1 |
| Simvastatin | 55 | 38 | 5 | 2 |
| Telmisartan | 20 | 8 | 1 | 1 |
| **TOTAL** | **515** | **433** | **78** | **92** |

**Supplementary Table 6 | Summary of $[M+H]^+$ Ions in the Feature-Based Molecular Network of the 12 Drugs.**

The table lists, for each drug, the precursor m/z, retention time (min), unique feature ID, component index (CI; representing the subnetwork ID within FBMN), and the number of nodes in each subnetwork. For Simvastatin, two features at m/z 419.2791 and 419.2792, both corresponding to the $[M+H]^+$ ion, were retained, as the compound eluted as two peaks (minor and major) at 4.75 and 5.01 min, respectively. Similarly, for Telmisartan, two features were included: the parent ion at m/z 515.2443 ($[M+H]^+$) and a fragment at m/z 258.1256 (library-annotated as 'Telmisartan'), both co-eluting at the same retention time.

| Drug | Precursor m/z | RT (mins) | ID | CI | Nodes |
|---|---|---|---|---|---|
| Cilnidipine | 493.1967 | 4.95 | 12221 | 5 | 98 |
| Clomifen | 406.1933 | 3.64 | 9988 | 129 | 42 |
| Erythromycin | 734.4685 | 2.39 | 6524 | 239 | 64 |
| Ketoconazole | 531.1563 | 2.63 | 7741 | 93 | 30 |
| Lansoprazole | 370.083 | 2.28 | 6063 | 63 | 5 |
| Loratadine | 383.1522 | 3.08 | 8893 | 373 | 14 |
| Metronidazole | 172.0717 | 0.58 | 2074 | 261 | 18 |
| Montelukast | 586.2179 | 5.14 | 12848 | 188 | 70 |
| Omeprazole | 346.1217 | 1.78 | 4975 | 1278 | 2 |
| Sertraline | 306.0812 | 2.83 | 8073 | 2 | 97 |
| Simvastatin | 419.2791, 419.2792 | 4.75, 5.01 | 11549, 12430 | 295 | 55 |
| Telmisartan | 515.2443, 258.1256 | 2.84 | 8143, 8175 | 208, 263 | 13, 7 |
